# Spatially-resolved Photoproximity Profiling of MYC Identifies a MYC-BAF Liability in Cancer Cells

**DOI:** 10.1101/2025.10.17.683176

**Authors:** Anthony J. Carlos, Shuyuan Huang, Dongbo Yang, Colin Swenson, Pratyasha Chakraborty, Shaopeng Yu, Benjamin D. Stein, Raymond E. Moellering

## Abstract

The c-MYC transcription factor is aberrantly expressed in most human cancers to enhance expression of proliferative gene programs. Owing to its pseudo-ordered structure and reliance on extensive and dynamic protein-protein interactions in distinct transcriptional regulatory complexes, defining context-specific MYC interactors has remained challenging. Therefore, mapping MYC-centered complex topologies in disease relevant models could identify components critical for its function which may serve as therapeutic targets in MYC-driven cancers. Here, we present a matched pair of photoproximity probes coupled with quantitative proteomics which enable context-dependent mapping of protein complex topology inside cells. We applied this spatially resolved, intracellular photoproximity (siPROX) profiling workflow to map MYC interactomes across temporal, spatial and disease-relevant contexts. Basal and inhibitor-treated profiles confirmed interactions with a wide range of known chromatin-associated transcriptional regulatory factors that define the extended MYC transcriptional bubble in live cells. Time-resolved mapping of inhibitor treated cells identified dynamic remodeling of numerous transcriptional regulatory factors and identified several BAF complex members (e.g., PBRM1 and SMARCC1)^1^ that persist in the presence of bromodomain inhibition. Furthermore, spatial MYC topology maps in small cell lung cancer cells confirmed the presence of BAF complex members under conditions where MYC induced target gene expression, altered cell morphology and enhanced proliferation. Lastly, loss of BAF function via inhibition of SMARCA2/4 ATPase activity resulted in rapid loss of chromatin-bound and nuclear MYC levels, downregulation of MYC-dependent transcripts and MYC-specific cell growth in several cancer cell models. Together, these data highlight the potential for siPROX to identify spatially resolved, dynamic TF interactors and highlight MYC-proximal BAF interactions as a targetable liability to regulate MYC-dependent transcription and proliferation.

## Introduction

Transcription factors (TFs) localize to gene enhancer and promoter regions of the genome to work in concert with chromatin remodeling complexes, epigenetic enzymes and basal transcriptional machinery to alter gene expression^2^. This enucleated cistrome works in concert with neighboring TFs, co-activating and co-repressive protein factors in a locus-, context- and time-dependent fashion. It is this dynamic assembly, disassembly and crosstalk that determines which genes are transcribed and to what extent entire gene programs are regulated. Therefore, the ability to map the membership within, and dynamic remodeling of, TF-centered protein complexes directly on chromatin *in situ* could provide an unprecedented view of gene regulation under both basal and dysregulated cellular states. Traditional approaches to identify TF-interaction partners have utilized co-immunoprecipitation (coIP) of either endogenous or transgenic TFs coupled to mass spectrometry for protein identification^3, 4^. While these approaches have been instrumental in the previous identification of many TF-interaction partners, there is significant potential for both false-negative and false-positive interactions due to the loss of sub-cellular compartmentalization and scaffolding afforded by chromatin architecture^4, 5^. Moreover, since transcriptional complex dynamics take place over timescales ranging from seconds to hours, it is difficult to capture transient dynamics that occur *in situ* with readouts that arise from equilibrated pull-downs hours or more later. For these reasons, developing general approaches to capture TF-centered protein complex dynamics directly in cells holds tremendous promise to accurately annotate the interaction landscape of the ∼1,600 TFs in the human proteome under (patho)-physiological contexts^2^.

c-MYC (referred to hereafter as MYC) is a prototypical and widely studied oncogenic basic-helix-loop-helix (bHLH) TF that binds to and activates genes broadly involved in driving cell proliferation^6, 7^. As a result, MYC is implicated as a master regulator of cellular physiology and one of the most frequently dysregulated oncogenes in human cancers^6–9^. Canonical MYC function and oncogenic potential requires heterodimerization with its obligate partner, MAX, enabling sequence-specific binding to E-box motif-containing enhancer and promoter elements for the activation of a broad network of target genes^10, 11^. A growing body of evidence suggests that MYC’s full range of function is mediated by a diverse and dynamic interaction landscape, encompassing both canonical and noncanonical protein-protein interactions (PPIs). A defining feature of MYC is its pseudo-ordered N-terminal transactivation domain (TAD), which enables the protein to engage in dynamic, multi-valent interactions with numerous cofactors involved in chromatin remodeling, transcriptional activation, and epigenetic regulation^12^ while maintaining faithful DNA binding capabilities with MAX. These include interactions with transcriptional coregulators such as TRRAP and GCN5, members of the larger STAGA complex^13, 14^, interactions with p300^15^, and interactions with components of the COMPASS-like methyltransferase complex such as WDR5^16^, thus allowing MYC to coordinate direct histone-modifying complexes. These interactions direct MYC and MAX localization to open chromatin to initiate and amplify proliferative gene programs.

The potential to map MYC-centered transcriptional complex topology and dynamics with high spatial precision directly in disease-relevant contexts could expand our understanding of MYC-dependent gene regulation and identify potential therapeutic paths to target MYC activity. Along these lines, intracellular proximity profiling methods like BioID^4, 17–19^ have recently been used to identify steady-state MYC interactors^17, 18^. However, the spatial and temporal resolution of BioID is intrinsically limited due to the low reactivity of the biotinyl-AMP anhydride tagging species^20^. In contrast, light-activated proximity profiling methods offer the potential for facile and non-perturbative delivery and activation of highly reactive – and therefore spatially restricted – labeling groups to profile direct protein interactors *in situ*^20, 21^. We previously reported a first-generation intracellular photoproximity methodology, PhotoPPI^22^, which utilized a modular chemical probe that could be delivered to any expressed protein of interest (POI) for in-cell, light-triggered proximity labeling and subsequent proteomic profiling of protein complex dynamics across rapid time scales. Subsequent photoproximity profiling methods have employed a combination of metal-centered photocatalyst/substrate pairs, which are predominantly restricted to cell-surface mapping due to poor membrane permeability and kinetics^23–25^, though recent examples have been reported with intein systems to deliver specific photocatalysts to the cell interior for proximity mapping^25^. Photocatalytic proximity methods recently expanded to employ catalyst/substrate pairs that allow for diverse radial mapping of cell surface complexes^26^ providing additional resolution beyond other proximity labeling approaches. Here, we sought to explore a suite of synthetic photoproximity probes that enable rapid, multiplexed and quantitative mapping of protein complex topology in live cells. We demonstrate that a pair of photochemically tuned, modular probes permit spatially-resolved, intracellular proximity (siPROX) profiling of basal MYC-centered complexes as well as dynamic changes in interactors in response to drug treatments and annotation in disease relevant models where MYC is dysregulated. In doing so, we identify novel MYC interactors and stable association of MYC pioneer complexes with BAF complex members. Due to the rapid and radial-tuning of multiplexed siPROX probes, we were able to map potential topological relationships between these members, as well as their dynamics in response to epigenetic remodeling drugs. Importantly, we show the therapeutic relevance of BAF complex activity on MYC stability, gene regulation and cell proliferation in MYC-induced oncogenic settings.

## Results

### Photochemically-tuned siPROX Probes Enable Proximal and Distal Target Protein Interaction Profiling

We previously reported a first generation photoproximity profiling (PhotoPPI) method using a modular, cell permeable small molecule probe, PP1^22^(Fig. 1A). Among the key features of this photoproximity probe was a central *o-*nitroveratryl group and a peripheral diazirine group for light-triggered activation and coincident diffusion of a reactive carbene for proximal protein tagging around a SNAP-tagged protein of interest (e.g., SNAP-POI) in cells (Fig. 1A). Importantly, the covalent labeling of the target protein permits facile delivery and washout of free probe prior to photoactivation, increasing profile specificity relative to methods with free substrate probe that can be activated by background photoactive species. The first-generation PhotoPPI system proved capable of in-cell proximity profiling of redox-sensitive and dynamic protein complexes, including the E3 ligase adaptor KEAP1. In theory, the fully synthetic nature of this system enables exploration of diverse photochemical groups to match and tune the central cleavage and reactive group activation properties and create a suite of probes with different effective diffusion and tagging radii (Fig. 1A). We reasoned that altering the steric properties on the benzyl carbon central to the nitroveratryl carbamate would alter the photocleavage of PP1-derivatives^27^. Indeed, a desmethyl derivative, AC1, showed slower photocleavage kinetics than PP1, whereas introduction of a bulky isopropyl group in AC3 accelerated photocleavage in bulk solution measurements (ED Fig. 1A-C). Incorporation of these derivative groups into modified photoprobes with identical diazirines resulted in a pair of probes: AC3 skewing toward faster relative cleavage before carbene activation for more rapid diffusion and distal protein labeling contrasted by AC1, which displays slower cleavage relative to carbene activation for more proximal protein labeling (Fig. 1A).

**Figure 1:**
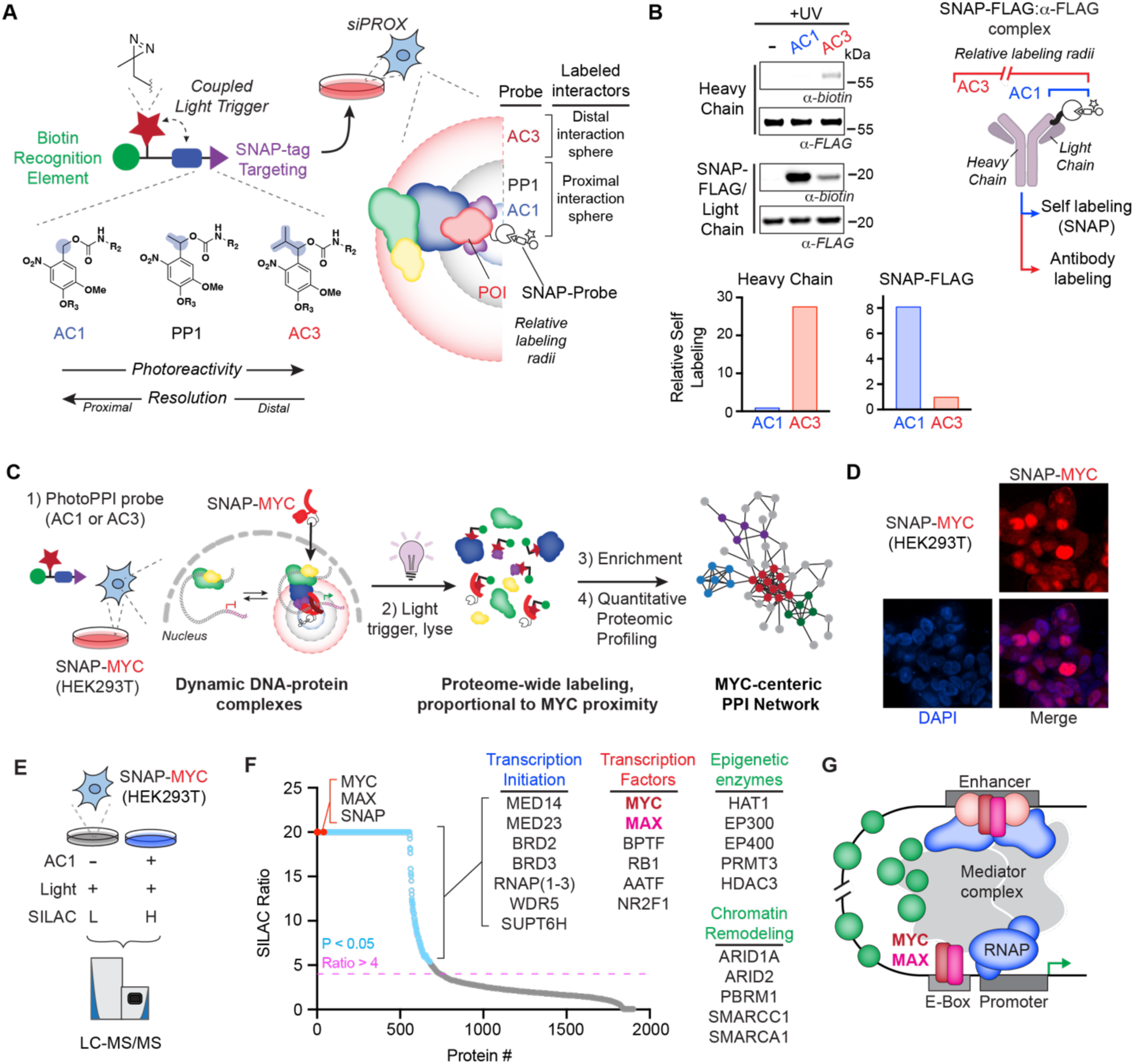
Photochemically tuned AC/AC3 probes enable spatially resolved mapping of protein complexes in vitro and in cells. **A)** Schematized structure of PhotoPPI probe. The photoactivatable coupled light trigger composed of a diazirine and tunable *o*-nitroveratryl (NV) linker is highlighted (left). Depiction of the relative theoretical labeling radii of the respective probes dependent on the reactivity of the NV linker (right). **B)** Differential AC1 or AC3 probe labeling of the SNAP-FLAG:α-FLAG complex. Immunoblots (α-biotin and α-FLAG) are representative of *in vitro* complex photolabeling by either AC1 or AC3 in SNAP-FLAG expressing cells. Bar plots show relative labeling of indicated complex members by each probe, as quantified by normalized intensity values. **C**) Schematic of Spatial-PhotoPPI (siPROX) workflow from live-cell probe treatment through bioinformatic analysis of MYC interactomes. **D**) Fluorescence microscopy confirming nuclear localization of the SNAP-MYC transgene in HEK293T cells. **E)** Experimental SILAC model using for precise Spatial-PhotoPPI mapping. **F**) Waterfall plot of enriched proteins showing SILAC ratios (AC1/DMSO) from HEK293T cells expressing SNAP-MYC. Enriched proteins were quantified in at least 2 biological replicates with a median SILAC ratio of 4 or greater. Highly enriched representative proteins are listed. Positive controls (MYC, SNAP and MAX) are highlighted in red. Enriched proteins with *p*-value < 0.05 are bordered in light blue; *n* = 4. **G**) Graphical depiction of MYC-centered super enhancer complexes as determined by siPROX.

Elaborated AC1 and AC3 probes containing these distinct photoreactive elements retained equivalent SNAP-tag protein labeling kinetics, potency, solubility and cell permeability (ED Fig. 3A-B). To interrogate the spatial labeling preferences of each probe, we performed on-bead proximity labeling with FLAG epitope-labeled SNAP-tag fusion proteins labeled in cells, then retrieved with bead-bound anti-FLAG IgG antibodies^22^. Covalent labeling with AC1 and AC3 followed by light activation at 365 nm resulted in significantly higher heavy chain protein labeling with AC3, consistent with the ability to diffuse further on average before carbene activation and protein labeling (Fig. 1B). By contrast, AC1 showed considerably higher self- and proximal light chain protein labeling. Experiments with a larger SNAP-Vinculin model protein showed the same spatial preference (ED Fig. 2). These labeling systems and photochemical constants are consistent with AC1 preferentially labeling a tight radius of ∼5-10 nm and AC3 targeting an extended and overlapping radius of ∼10-30 nm^23, 28^.

To interrogate MYC interactors in cells, we first stably expressed a SNAP-MYC fusion protein in HEK293T cells (Fig. 1C-D). To confirm similar expression, localization and regulation of the transgene, we utilized the SNAP tag for imaging and turnover measurements. Immunofluorescence imaging and immunoblot confirmed predominantly nuclear localization of SNAP-MYC at levels ∼8-fold above endogenous MYC levels in parent HEK293T cells (Fig. 1D ED Fig. 9A). Pulse-chase measurements with AC1 probe showed that SNAP-MYC had a half-life of approximately 50 min in cells, in line with previously published stability measurements for endogenous MYC (ED. Fig. 3D). Interrogation of other stable (e.g., LDHA) and transient (e.g., NRF2) SNAP-tagged proteins also showed expected half-lives, confirming that neither SNAP tag fusion or AC1/3-probe labeling perturbs general protein localization or stability (ED Fig. 3C).

As in our first-generation PhotoPPI workflow, we utilized stable isotope labeled cells (stable isotope labeling of amino acids in cell culture, SILAC) expressing SNAP-MYC to generate a basal MYC interactome in live cells. AC1- or vehicle-treated SILAC cells were irradiated with light for 5 min, followed by lysis and enrichment of biotin-tagged proteins for quantitative LC-MS/MS proteomics (Fig. 1E). SILAC quantification of AC1-enriched targets confirmed robust enrichment of a MYC-centered interaction network under these conditions (Fig. 1F-G). Notably, MYC and its obligate dimerization partner MAX, alongside the SNAP labeling protein itself (an engineered version of MGMT) were enriched at the top of the profile with SILAC ratios of >20. Many known MYC interacting proteins were enriched by AC1 in these cells, including epigenetic co-activators like EP300, EP400, HAT1, PRMT3, WDR5 and HDAC3 (Fig. 1F-G). Members of transcription initiation complexes and core RNA polymerases 1-3 were enriched, alongside enhancer-associated proteins known to associate with MYC like BRD2, BRD3, MED14, TRRAP and MED23 (Fig. 1F-G). Numerous chromatin remodeling enzymes and members of the BAF complexes like ARID1A, ARID2, PBRM1, SMARCC1 and SMARCA1 were also observed in proximity to MYC^29, 30^. Finally, several other general and specific transcription factors (BPTF, RB1, AATF, NR2F1 and RELA) as well as proteins associated with DNA damage, regulation of chromosome organization, cell cycle control and mitotic progression were enriched proximal to MYC-centered proximity labeling in cells (ED Fig. 5E-H). Importantly, we performed the same siPROX experiment on SNAP-tagged Vinculin, which is an unrelated, cytosolic structural protein and observed little overlap in the proximity profiles of these two (ED Fig. 4). Collectively, this basal siPROX profile paints a high-resolution map of both known and novel MYC-associated protein complex members directly in live cells. A comparison of published MYC interactors derived from BioID profiling^17^ and MYC interactors derived from siPROX profiling showed a stark increase in identified MYC interactors using siPROX. Following Metascape analysis on both lists, we observe strong overlap at the pathway level with shared pathway terms related to cell cycle, MYC targets and SWI/SNF superfamily-type complex. It should be noted that BioID MYC interactomes were acquired in the presence of tetracycline, proteasome inhibitors and high concentrations of biotin over a 24 hr labeling window This stands in stark contrast with the rapid (∼5 min) and minimally perturbative siPROX profiling workflow.

### Time-Resolved Mapping of MYC Complex Dynamics in Response to Epigenetic Drug Treatment

We hypothesized that the facile probe delivery and covalent labeling possible with siPROX probes followed by rapid light activation lends itself to profiling dynamic changes to protein complex topology in response to altered signaling and/or drug treatments. To test this premise, we performed quantitative comparisons of MYC-centered siPROX profiles in response to bromodomain-containing reader protein inhibition with the small molecule JQ1^31^ across minutes-to-hours timeframes (Fig. 2A; ED Fig. 5A). We first tested the effect of extended (24 hr) JQ1 treatment on MYC-proximal protein profiles in SNAP-MYC-expressing HEK293T cells. Many of the enriched interactors detected in the AC1 vs. Veh. basal profiles were detected here (ED Fig. 5B-C), with the added quantitative comparison of DMSO vs. JQ1 treatment to directly quantify changes in MYC interactors agnostically across the proteome (Fig. 2A). Probe self-labeling on SNAP-MYC and core interactors like MAX were not significantly altered with JQ1 treatment (Fig. 2C-D). By contrast, many of the extended transcriptional regulatory protein network associated with MYC was significantly altered (Fig. 2A-B, ED Fig. 5C-F). This included loss of BRD2 and BRD3 interactions, which are expected as they are known targets of JQ1, alongside RNA polymerase (e.g., POLR3A), epigenetic enzymes like EP300, EP400 and HDAC3. Other proteins involved in DNA repair and cell cycle regulation were also reduced upon extended bromodomain inhibitor treatment. Intriguingly, several bromodomain-containing proteins remained in proximity to MYC, including the BAF complex member PBRM1 and its co-complex member SMARCC1 (Fig. 2C, ED Fig. 5E).

**Figure 2:**
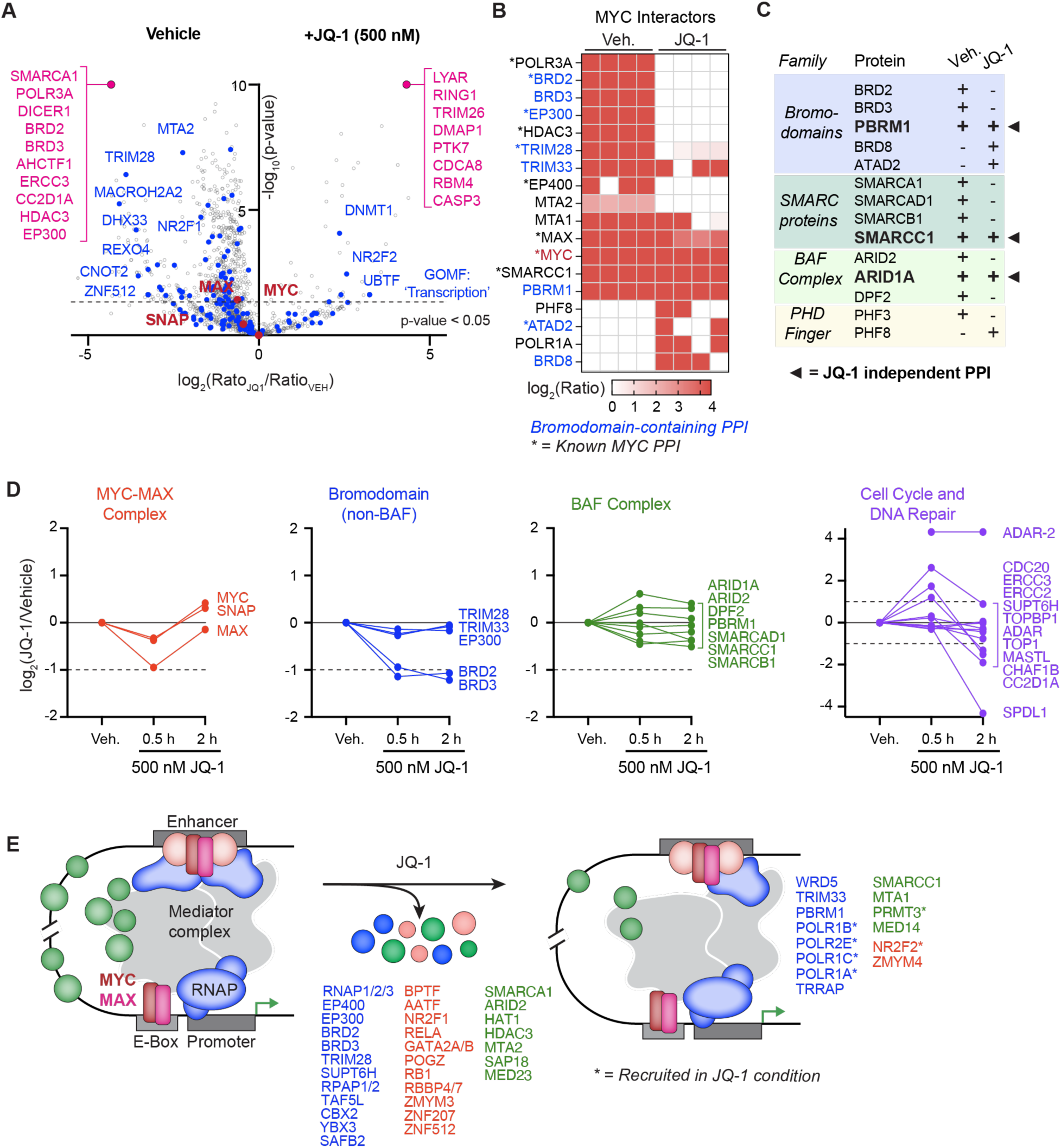
Mapping the dynamic MYC interactome in response to bromodomain inhibition reveals JQ-1 resistant MYC-BAF interactions. **A**) Volcano plot comparing SILAC ratios derived from HEK293T cells expressing SNAP-MYC under basal (left) and JQ-1 treatment (500 nM; right) conditions; n = 4 biological replicates. **B**) Heatmap of select chromatin associated (GO) proteins plotted in A. **C**) Tabulated bromodomain, BAF and other chromatin regulatory MYC-interacting proteins enriched in basal and JQ1 treated cells. **D**) Dynamic shifts in the MYC interactome following JQ-1 treatment. Red: control SNAP-MYC/MAX complex; blue: bromodomain proteins; green: BAF complex members; purple: cell cycle and DNA repair proteins. **E**) Graphical depiction of dynamic MYC-chromatin regulatory hubs before (left) and after (right) JQ-1 treatment. Asterisks denote interactors present only after JQ-1 treatment.

To probe more acute changes to MYC-associated complexes, we performed similar siPROX experiments at 30 mins and 2 hrs of drug treatment. Multiplexing across short kinetic windows like these are enabled by the streamlined siPROX workflow and rapid kinetics of probe activation with minutes of light exposure. As with the basal and extended JQ1 treatment datasets, the 0.5 and 2 hr profiles contained many members of the core and extended MYC transcriptional complex (ED Fig. 6, Supplemental Data Table 1). Proximity labeling of the core SNAP-MYC/MAX complex remained largely unaffected, which contrasts with rapid and significant loss of BRD2 and BRD3 association with MYC at both 0.5 and 2 hr timepoints (Fig. 2D). Exemplary co-activator and core transcriptional machinery proteins like EP300, TRIM28, MED14 and POL3RA remained stably associated with MYC at these acute timepoints, contrasting with their widespread reduction in proximity after extended JQ1 treatment (Fig. 2B-D). Clusters of proteins associated with cell cycle progression and DNA repair, including TOP1, MASTL, SPDL1 and others showed significant reduction in MYC proximity between the 0.5 and 2 hr timepoints. As in the longer JQ1 treatment dataset, a large number of BAF complex members were found proximal to MYC and were unperturbed upon acute drug treatment. This is intriguing considering that several of these proteins, for example PBRM1, contain bromodomains that are involved in chromatin localization, offering the potential that MYC-BAF association is mediated by contacts beyond general BAF-chromatin interactions at actively transcribed loci. These data collectively paint a picture of rapid and sustained loss of the core JQ1 targets BRD2 and BRD3 in MYC complexes in as little as 30 minutes, followed by more widespread reduction in MYC complex associations with proteins involved in DNA repair and cell cycle progression (Fig. 2D-E). Upon sustained inhibition the MYC-associated interactome is broadly rewired with sustained reductions in MYC proximal co-activators, transcription factors, DNA repair and core transcriptional machinery (Fig. 2E). These MYC-centered siPROX profiles thus provide high resolution snapshots of MYC-associated complexes on chromatin consistent with the known importance of specific bromodomain-containing proteins for MYC-dependent transcription.

### Multiplexed siPROX Identifies Proximal and Distal MYC Complex Topology in MYC-Dependent and Pathologically-Transformed Cancer Cells

Basal and inhibitor-treated MYC-centered profiles in a model cell line (HEK293T) confirmed the ability to identify and track known and novel MYC interactors in cells with siPROX probes. To identify and specifically map the spatial relationships of these complex members in cancer-relevant models, we established and applied a multiplexed and quantitative siPROX workflow using AC1 (proximal-focused) and AC3 (distal-focused) in the same experiment. We used a triple-SILAC labeling scheme to compare vehicle, AC1 and AC3-enriched proteins in the same siPROX experiments owing the precision of metabolic-isotope labeling, streamlined workflow and minimal sample loss. We first applied multiplexed siPROX to the same SNAP-MYC expressing HEK293T cells and compared AC1-biased to AC3-biased interactors. MYC, MAX and SNAP protein enrichment was biased toward AC1 relative to AC3 (Fig. 3A-B), which is consistent with *in vitro* experiments (Fig. 1B) and confirms the theoretical photochemical tuning of siPROX probes translates to live cells. Globally, there is significant overlap in the profiles of AC1 relative to AC3, however AC3 enriched roughly 3-fold more total proteins. Some members of the core transcriptional machinery like POLR2A are biased in the AC1 profile, whereas most chromatin remodeling proteins, components of the mediator complex, epigenetic modifiers and transcription factors were more highly enriched in the distal-AC3 siPROX profile (Fig. 3B). The AC3-biased profile also contained approximately 5-fold more highly enriched targets relative to AC1, which is indicative of proteins that are likely present at lower frequency in proximity of MYC or are longer distances from MYC (ED Fig. 8C-F).

**Figure 3:**
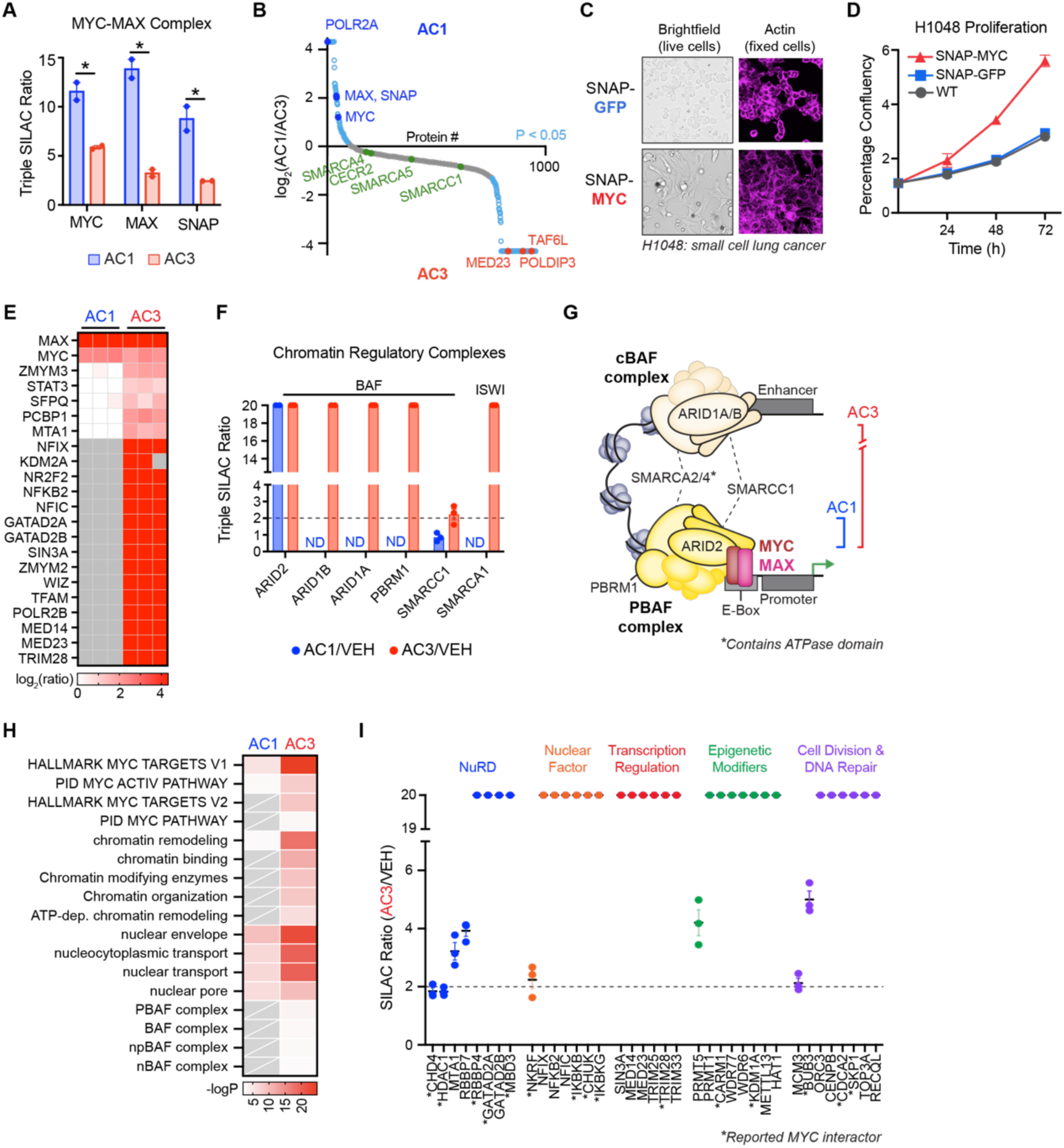
Overexpression of MYC in small-cell lung cancer leads to profound phenotypic changes accompanied by MYC-BAF crosstalk. **A**) siPROX analysis showing Triple SILAC ratios for the MYC-MAX complex. **B**) Waterfall plot of log_2_(AC1/AC3 ratio) against protein number. Highlighted proteins are preferentially labelled by either AC1 (blue) or AC3 (red). Proteins labelled by both probes are highlighted in green. **C**) Representative live-cell brightfield (left) and fixed-cell actin (phalloidin stain; right) images of small cell lung cancer (H1048) cells stably expressing SNAP-MYC or -GFP. **D**) Relative proliferation of SNAP-MYC (red), -GFP (blue) and WT (grey) H1048 cells, as assessed via CellTiter-Glo (CTG) assay. **E-F**) Multiplexed siPROX profiling of SNAP-MYC H1048 cells. Heatmap indicates log**_2_**(Probe ratio) for known and novel MYC interactors (E). Bar plot shows log_2_(Probe ratio) for known and novel MYC interactors (F). **G**) Proposed model of differential MYC membership across cBAF and PBAF complexes. **H**) Metascape analysis of proteins preferentially enriched by AC1 and AC3. **I**) Scatterplot reporting SILAC ratios (AC3/Vehicle) for MYC-interacting protein clusters and complexes. n = 2 (A-B) and 3 (E, F, H, I) biological replicates.

Next, we were interested to use multiplexed siPROX to determine whether and to what degree the MYC-centered complexes detected in HEK293T cells were present in more physiologically relevant tumor cell settings. We focused on small cell lung cancer (SCLC), where many neuroendocrine-like tumors are known to harbor high MYC expression and MYC-dependence in cell, animal and patient-derived tumor models^32–34^. We expressed SNAP-MYC in the NCI-H1048 cancer cell line and confirmed expression of the transgene at ∼4-fold higher levels than the endogenous protein, alongside compensatory reduction in endogenous MYC expression (ED Fig. 9A). Turnover of SNAP-MYC was similar to that in HEK293T cells and other previously published settings (ED Fig. 3D), collectively establishing a model where MYC is expressed and regulated within a range that is commonly found in SCLC cell lines and tumors^35, 36^. Intriguingly, the morphology of SNAP-MYC expressing H1048 cells changed drastically relative to parental or SNAP-GFP expressing control cells, becoming significantly larger with pronounced filopodia and mesenchymal character (Fig. 3C). SNAP-MYC expressing cells also harbored increased expression of known MYC target genes (Fig. 4A) and showed enhanced proliferation relative to SNAP-GFP or parental control cells (Fig. 3D).

**Figure 4:**
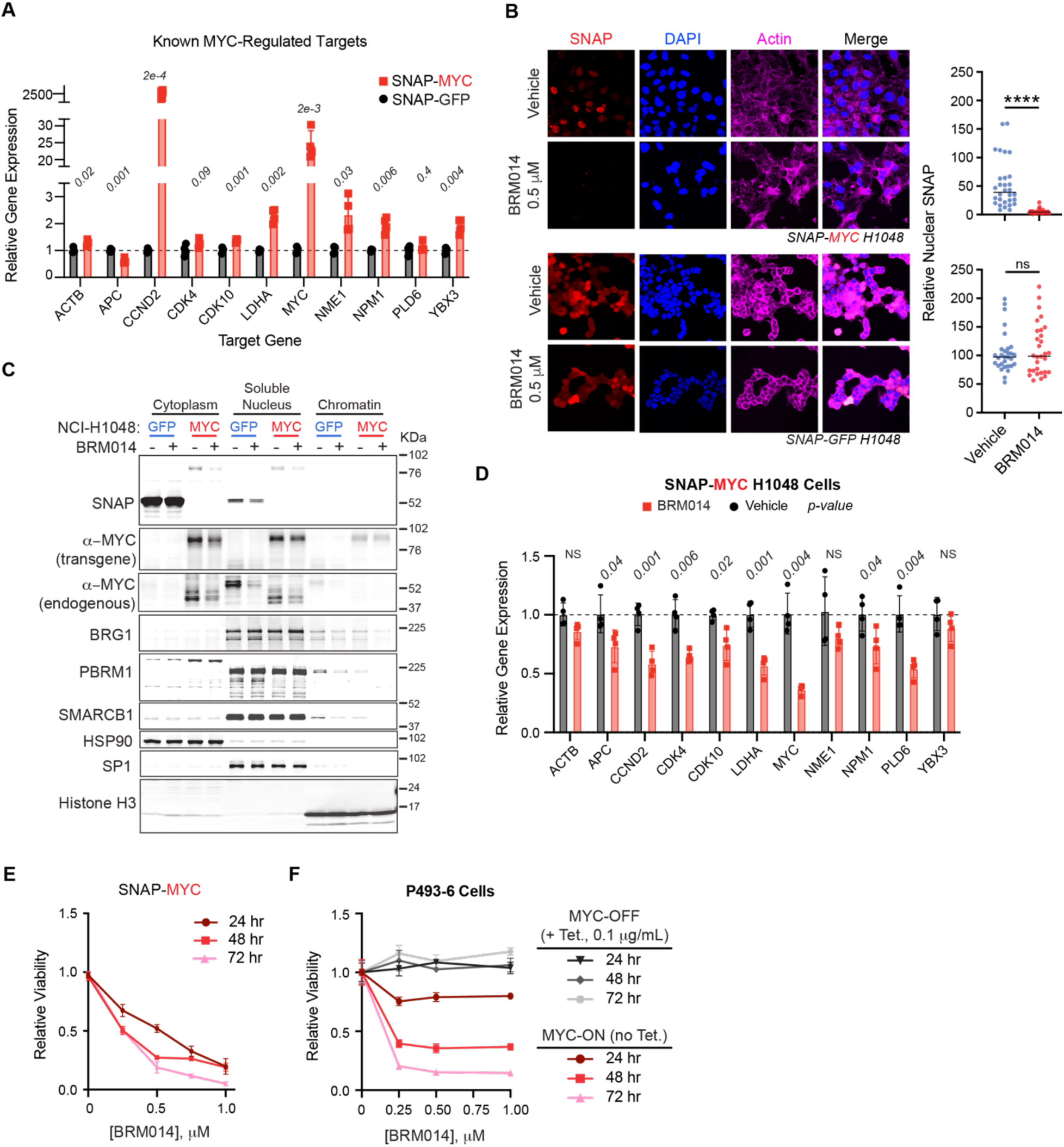
BAF complex inhibition reduces MYC stability, signaling and proliferation in models of MYC-driven cancer. **A**) Relative qPCR on MYC target genes in H1048 cells expressing SNAP-MYC or -GFP; n = 4. **B**) Representative confocal images of SNAP-MYC and -GFP H1048 cells treated with BRM014 inhibitor (500 nM, 18 h). Quantification shows relative levels of nuclear SNAP-tagged MYC (top) or GFP (bottom). **C**) Effect of BRM014 treatment on levels of SNAP, MYC, BAF complex members (BRG1, PBRM1, and SMARCB1) levels in the nucleus, cytosol and chromatin. HSP90, SP1, and Histone H3 were used as controls for fractionation. **D**) qPCR analysis of MYC target genes in response to BRM014 treatment in SNAP-MYC H1048 cells. **E**) Relative viability of SNAP-MYC H1048 cells following treatment with indicated doses on BRM014. **F**) Relative viability of proliferating P493-6 cells starting in either MYC-ON (reds, pink) or MYC-OFF (black, grays) state and treated with indicated doses of BRM014.

We performed multiplexed siPROX profiling in SNAP-MYC H1048 cells and confirmed general enrichment of the core MYC/MAX complex and an extended interaction network similar to that observed in HEK293T cells (Fig. 3E-G, ED Fig. 9C-D). As in HEK293T cells, the siPROX profile was significantly biased toward AC3-enriched distal interactors. These protein targets included conserved members of the core transcriptional machinery (POLR3A), co-activators (EP300, BRD2, WDR5), DNA replication and repair enzymes (TOP3A, RECQL, ORC3, MCM3), transcription factors and regulators (NFKB2, NFIC, NFIX, NKRF) and cell cycle control proteins (CDCA2, SKP1) (Figure 3H-I). In this context, MYC was heavily linked with NFkB pathway members, including NFKB2, NKRF, CHUK, IKBKB, IKBKG and others, consistent with known connections between reciprocal regulation of MYC and NFkB-dependent signaling in cancer^37–39^. BAF complex members, including ARID1A & B, PBRM1, SMARCA1 were also highly enriched in the extended AC3-biased profile alongside ARID2 and SMARCC1 that were detected by both AC1 and AC3, suggesting closer proximity in MYC complexes (Fig. 3E-F). Notably, GO analyses confirmed enhanced enrichment of pathways related to MYC pathways, chromatin remodeling, nuclear transport and BAF complexes by AC3 in MYC-expressing H1048 cells (Fig. 3H). In addition, comparative pathway analysis of HEK293T and H1048 siPROX enriched proteins revealed common and distinct features. Both cell lines harbored common mapped pathways including core MYC pathway members, cell division, mitotic progression, DNA repair and transcription factor binding (ED Fig. 9B-C). While the specific membership and spatial topology of larger co-regulatory complexes likely change in different cell settings, these data confirm the ability of siPROX to identify and differentiate highly proximal and more distal members of MYC-centered complexes directly on native chromatin in live cells. Moreover, these siPROX profiles establish that core members of the BAF complex are in stable and extended MYC transcriptional complexes, implicating potential MYC-BAF functional interactions in contexts where MYC drives pathological phenotypes (Fig. 3F-G).

### BAF Protein Activity Regulates MYC Stability and Oncogenic Phenotypes in MYC-Driven Cancer Cells

BAF complex activity has been associated with MYC function in diverse settings, including CD8^+^ T-cell maturation^40^ and in SCLC and multiple myeloma studies surrounding POU2F3+ SCLC and IRF4 dependencies, respectively. Specifically PROTAC degradation of BRG1/BRM1 led to observed decreases in cMyc expression in SCLC cell lines, specifically those dependent on POU2F3 expression^41–43^. Combined with our siPROX profiles here that place core MYC and BAF complexes in high proximity in cancer cells, we hypothesized that targeting BAF complex activity could directly impact MYC function. Specifically, the stability of these complexes, including bromodomain-containing members like PBRM, in the presence of bromodomain inhibitor treatment led us to hypothesize that BAF complexes may participate in recruitment of MYC to and stabilization on sites in open chromatin. To test this premise, we used the published compound BRM014, which targets the ATP binding site of SMARCA2/BRM^44^ and has been shown to regulate BAF chromatin remodeling activity in cells and animals. Visualization of SNAP-MYC protein in H1048 cells demonstrated that BRM014 (500 nM) treatment significantly reduced nuclear-localized and total MYC protein levels in cells (Fig. 4B). Sub-cellular fractionation further supported this effect, as nuclear and cytosolic SNAP-MYC levels were reduced with BRM014 without significant perturbation of BAF protein levels or localization (e.g., BRG1, PBRM and SMARCB1; Fig. 4C). By contrast, SNAP-GFP levels were unaffected by BRM014 by imaging and sub-cellular fractionation (Fig. 4B-C and ED Fig. 10A). BRM014 treatment significantly reduced MYC-dependent transcripts involved in cell cycle regulation, metabolism, protein synthesis and RNA processing in both SNAP-MYC-expressing and parental H1048 cells (Fig. 4D; ED Fig.10B). This effect was followed by significant time- and dose-dependent anti-proliferative activity in SNAP-MYC expressing cells (Fig. 4E), as well as SNAP-GFP expressing H1048 cells that also express and rely upon MYC. To determine whether the effect of BRM014 treatment on proliferation is MYC-dependent, we tested its activity in P493-6 cells, which contain exogenous MYC expression under control of tetracycline. BRM014 led to significant dose- and time-dependent anti-proliferative activity of ‘MYC-ON’ cells and essentially no effect on ‘MYC-OFF’ cells, where MYC expression was ablated by tetracycline treatment (Fig. 4F). Collectively, these data confirm that targeting BAF complex activity reduces MYC-dependent signaling and proliferation in cancer cells of diverse origin.

## Discussion

Here we describe a general approach to map intracellular protein complex dynamics and spatial relationships using siPROX chemical probe multiplexing and a quantitative proteomics workflow. While the modularity of siPROX should be applicable to interrogate essentially any intracellular protein complex, we first applied it to profile a TF, and MYC in particular, due to the importance of context-specific and dynamic protein interactions for its role in transcriptional regulation and oncogenic potential. We demonstrate that siPROX profiling of MYC-centered complexes captures many known interactors spanning obligate binding partners like MAX and associated co-activators, core transcriptional machinery and extended members of the MYC transcriptional bubble, redefining the MYC cistrome. Essentially all previously established MYC interactors identified from many studies were captured simultaneously and in an unbiased fashion in a single siPROX experiment, underscoring the potential for discovery and quantitative complex mapping inherent to siPROX and proximity profiling methods in general. A unique aspect of our siPROX system is the use of direct photoactivatable probes delivered via a biorthogonal, covalent anchor to a protein of interest (i.e., SNAP-MYC). This builds on our first-generation PhotoPPI system and contrasts with other predominantly photocatalytic approaches. We demonstrate here that direct siPROX probe delivery enables rapid and minimally perturbative labeling and photoactivation kinetics to yield robust labeling of MYC-centered protein complexes in live cells. Therefore, despite the loss of presumed signal enhancement with a catalytic system, we believe that local photoactivation only at the target of interest with siPROX can produce highly specific and precise profiles for SNAP-MYC or other proteins of interest in the future. We also believe our dataset underscores the power of photoactivation in contrast to much less-reactive and more perturbative biorthogonal methods like BioID and APEX. Indeed, the one previous published proximity profile of MYC interactors utilized a 24 hr labeling time course with BioID in the presence of proteasome inhibition, which we posit is incompatible with a short half-life protein like MYC or for mapping of dynamic complex rewiring. Our comparison with this dataset shows modest overlap, a lack of many known interactors captured by BioID, coupled with much higher coverage of known and novel interactors using siPROX. Recent advances in the kinetics and chemistry of other enzyme-based systems could improve in these areas^45, 46^ and synergize with siPROX. Collectively, siPROX presents a high precision, biorthogonal platform self-contained within a multiplexed set of probes portable to any protein of interest relative to other systems that rely on dual probe activation or extensive protein engineering^25^.

Beyond mapping basal interactomes, we show that multiplexed siPROX takes advantage of carefully tuned photokinetics within AC1 and AC3 to preferentially capture interactors that are more proximal and distal to the target of interest, respectively. We confirmed this both *in vitro* and across several datasets in diverse cellular settings. Indeed, we showed a strong preference for AC3 to capture a larger number of protein interactors in the extended MYC cistrome, including several co-activator proteins, chromatin remodeling complexes and mediator proteins. By contrast, AC1 strongly favored proteins like MAX, that are known to be highly proximal to MYC. Thus, these probes can be used in tandem to map a spatial continuum around a target of interest. A second and related capability of siPROX is the potential to track dynamic changes to MYC-centered complexes along physiologic kinetic regimes (i.e., minutes-to-hours). We demonstrated this feature through tracking the dynamics of the MYC interaction landscape topology in response to the bromodomain inhibitor, JQ1, which identified rapid dissociation of canonical targets like BRD2 and BRD3, followed by a deeper and more sustained change in MYC association with co-activators, general transcriptional machinery and chromatin remodeling complexes.

Finally, the capacity to profile bromodomain inhibitor response in spatially-resolved MYC-centered complexes enabled the identification of stable and persistent MYC association with an extended network of BAF complex proteins, with the novel observation of a persistent pBAF specific complex member, PBRM1. We were further intrigued by the distal association of cBAF and NuRD chromatin remodeling complexes alongside proximal association with pBAF complex member PBRM1, even in the presence of extended bromodomain inhibitor treatments (Fig. 2B-C). These collective profiles suggested that MYC-BAF association may be involved in chromatin localization, stability and transcriptional activity of MYC in cancer cells, which extends previously published connections between BAF activity and MYC function in T-cells^40^ and MYC interactions with other chromatin remodelers in distinct cancers^47^. We found that pharmacologic inhibition of SMARCA2/SMARCA4 activity resulted in significant reduction of MYC protein levels and association with chromatin in H1048 cells, a POU2F3 positive SCLC cell line previously shown to be dependent on BAF function^41, 42^. Indeed, constitutive SNAP-MYC overexpression caused pronounced morphological transformation, enhanced MYC transcription and increased cellular proliferation. Allosteric inhibition of BAF ATPase activity^44^ reduced MYC-dependent target gene expression and growth in these cells, as well as a MYC-dependent Burkitt’s lymphoma cell line, P493-6. Additional studies are needed to mechanistically explore this connection and potential for therapeutic targeting of MYC-dependent tumors. Collectively, siPROX significantly expands upon the resolution of intracellular photoproximity profiling strategies for precise detection and relative quantification of proximal and distal protein complex membership, here demonstrated around a master oncogenic TF. Future application of this approach may be powerful for understanding small molecule drug mechanism of action, study of fundamental transcriptional dynamics and therapeutic target discovery.

## Methods

### General synthetic methods

Please Supplemental Methods for synthetic methods and characterization of siPROX chemical probes.

### Cell culture

HEK293T and H1048 cell lines were acquired from ATCC. HEK293T cells were propagated in DMEM (Corning) supplemented with 10% fetal bovine serum (FBS, Corning) and 1% penicillin/streptomycin (Gibco). All cell lines were grown at 37 °C in a 5% CO_2_ humidified incubator. All cell lines tested negative for mycoplasma using the Lonza MycoAlert PLUS Detection Kit.

### Cloning and Fusion

Generation of N-terminal 3xFLAG-SNAP-tag destination vector:

All PCR reactions were performed using NEB Q5 high-fidelity polymerase (M0491S) and NEB dNTP solution mix (N0447L). SNAP tag fragment was generated from pSNAP_f_ Vector (NEB #N9183S) with primer F1 and R1 and incorporated into linearized vector pDEST_N_FLAG_HA_IRES_puro (Addgene #41033) with primer F2 and R2.

F1: GAGGTTGATCTGCCACCATGGACTACAAAGACCATGACG

R1: GTACAAACTTGTTTGATAACCCAGCCCAGGCTTG

F2: CAAGCCTGGGCTGGGTTATCAAACAAGTTTGTACAAAAAAG

R2: GTCTTTGTAGTCCATGGTGGC

pNTerm-3xFLAG-SNAP-MYC:

pDONR223 MYC was acquired from Addgene (Plasmid #82927). The plasmid was amplified in DH5alpha in the presence of Spectinomycin. pDONR223 MYC was cloned into pNTerm-3xFLAG-SNAPtag destination vector using LR Clonase (NEB) following manufacturer protocol.

pNTerm-3xFLAG-SNAP-GFP:

Ligation reaction was performed using Gateway LR Clonase (Invitrogen #11791100.) pDONR223 GFP was obtained from Addgene (#209057.) pDONR223 GFP was cloned into pNTerm-3xFLAG-SNAPtag destination vector using LR Clonase following manufacturer protocol.

pNTerm-3xFLAG-SNAP-NRF2:

Ligation reaction was performed using Gateway LR Clonase (Invitrogen #11791100.) pDONR223 NFE2L2 was obtained from Addgene (#81259.) pDONR223 NFE2L2 was cloned into pNTerm-3xFLAG-SNAPtag destination vector using LR Clonase following manufacturer protocol.

### Stable SNAP-POI Cell Line Generation

Mammalian cells stably expressing target protein constructs were obtained by transfecting a 6 cm plate of HEK293T cells with 1.5 ug Gag-Pol (Addgene #12251), 1 µg VSV-G (Addgene #12259), 0.5 µg Rev (Addgene #12253), and 2 µg of plasmid of interest with 12.5 μL of lipofectamine 2000 (Invitrogen #11668019) in Opti-MEM (Gibco #31985070) for 4 hours. The media was replaced with DMEM (Corning #MT10017CM) supplemented with 10% FBS (Corning MT35010CV) and 1% pen (Gibco # 15140122); resultant viral media was collected in 48 hours, passed through a 0.45-micron filter, and supplemented with 8 µg/mL polybrene (Sigma). Viral transduction was achieved by culturing a separate population of mammalian cells in the diluted viral media for 24 hours. Afterward, the viral media was removed and the transduced cells grown in full media containing 2.5 µg/mL puromycin (Gibco #A1113803). Stable incorporation of the transgene was confirmed by western blotting using monoclonal anti-FLAG M2 antibody (Sigma).

### SDS-PAGE and Immunoblot

Cells were lysed by on-well lysis using ice-cold 1 x RIPA (Millipore) and tip sonicated (Fisher Scientific FB-505) over ice. Insoluble debris was cleared by centrifugation, and the supernatant was diluted into 4X Laemmli buffer containing 50 mM dithiothreitol (DTT) as a reducing agent. Samples were prepared for SDS-PAGE by heating to 95 °C for 5 min, cooled to room temperature, resolved on 10% SDS-PAGE gel, and transferred to nitrocellulose membranes by standard western blotting methods. Membranes were blocked in 2% BSA in TBS containing 0.1% tween-20 (TBST) and probed with streptavidin-800 (Licor Odyssey CLx) to visualize biotin labeling. Blot intensities were quantified in Image J and normalized in Microsoft Excel.

### *In vitro* photo-kinetics experiments

A 200 μM solution of model nitroveratryl carbamates in methanol was prepared from 10 mM DMSO stock solutions. 1 mL of the solution was added to three individual Eppendorf tubes for three 3 replicates. The samples were placed on ice and positioned approximately 4 cm the source of irradiation. The samples were irradiated with 365 nm light using a Spectrolinker XL-1500a (Spectroline) for several time points and analyzed via LC-MS on an Agilent 1200 Series G1311A using the following method [Buffer A: 95% H2O, 5% MeCN, .1% TFA. Buffer B: 95% MeCN, 5% H2O, 0.1% TFA. 0 min (85% A, 15% B), 2min (85% A, 15% B), 2.5 min (60% A, 40% B), 8 min (0% A, 100% B), 9.75 min (0% A, 100% B), 10 min (85% A, 15% B), 12 min (85% A, 15% B) at .5 mL/min. Integration data for UV (215 nm) were used to derive kinetic parameters on Graph Pad Prism 8.

### *In vitro* α-FLAG photolabeling experiments

HEK293T cells with stable transduction of SNAP-Flag, SNAP-Flag-GFP, or SNAP-Flag-VCL were collected and lysed in 1xRIPA w/ 1mM DTT supplemented with cOmplete protease inhibitor (Roche #04693159001) and clarified. Cell lysates were normalized to 1mg/mL in 1.5mL, mixed with 240uL of washed anti-FLAG M2 affinity gel (Sigma A2220) and rotated at 4 °C overnight. Samples were then spun down at 1500 rpm, supernatant aspirated, and the resin washed with DPBS supplemented with 1mM DTT (1 mL x 3). The washed lysate-resin suspension was partitioned and incubated with vehicle or 500 nM of probe for 1 hour at 4 °C in dark on rotation. Samples were spun down at 1500 rpm, supernatant aspirated, and the resin washed with DPBS supplemented with 1mM DTT (1 mL x 3). The samples were then irradiated with 365nm light for 5 min. Samples were spun down, supernatant aspirated, and the resin washed with DPBS. The resin was boiled 95 °C in 20 µL 4x-loading buffer containing 8% SDS and 400 mM DTT. After which, 60 µL of DPBS was added and the suspension boiled at 95 °C for an additional 5 minutes. The samples were then run on SDS-PAGE gel and transferred to nitrocellulose where they were stained with anti-mouse IR dye (Li-cor #926-68072) and streptavidin IR dye (Li-cor #926-32230). Bands corresponding to biotinylated proteins and the FLAG antibody were visualized using an Odyssey infrared imager (Li-cor).

### PhotoPPI probe dosing on live cells for immunoblot analysis

HEK293T cells expressing SNAP-tagged proteins of interest were seeded in 24-well plates at 100,000 cells per well in 1.0 mL of RPMI media (Corning) supplemented with 10% fetal bovine serum (Corning) and 1% penicillin/streptomycin (Gibco) 24 hours before the experiment. Cells were treated with 0, .5, 1, 5, 10 and 25 μM of AC1 in 500 μL of serum-free, phenol-red-free RPMI media (Gibco) for 1 h at 37 °C. The cells were then washed in 1 mL of PBS twice and lysed in ice-cold 1x RIPA buffer supplemented with protease inhibitor (Roche). Cells were lysed while shaking on ice for 20 min before transfer and tip sonication (3 x 1 s pulses at 30% amplitude). The lysates were collected and processed for anti-biotin immunoblot analysis.

### Pulse Chase Experiment on live cells

HEK293T cells expressing SNAP-tagged proteins of interest were seeded in 24-well plates at 100,000 cells per well in 1.0 mL of RPMI media (Corning) supplemented with 10% fetal bovine serum (Corning) and 1% penicillin/streptomycin (Gibco) 24 hours before the experiment. Cells were pre-treated with no treatment/10 μM TBHQ/2.5 μM MG132 for 1 hr at 37°C. The cells were then treated with 20 μM of AC1 in 0.25 mL serum-free RPMI media with corresponding activator treatment and were incubated for 1 hr at 37°C. The cells were washed in 0.25 mL serum-free RPMI media with corresponding activator treatment for 10 min. The cells were incubated in full media for 0-12 hour and were collected with RIPA buffer supplemented with protease inhibitor (Roche). The lysates were collected and processed for anti-biotin immunoblot analysis.

### SNAP-MYC localization experiment using HEK293T cells

384-well plates (Greiner) were pretreated with 0.01% poly-L-lysine (Sigma-Aldrich) for 10 minutes and dried for 2 hours at room temperature. HEK293T cells expressing SNAP-tagged proteins of interest were seeded in 384-well plates at 2,500 cells per well in 50 μL of RPMI media (Corning) supplemented with 10 % fetal bovine serum (Corning) and 1% penicillin/streptomycin (Gibco) 48 hours before the experiment. The cells were then treated with 3 μM of SNAP-cell TMR-STAR (NEB #S9105S) in 50uL of full media and incubated for 0.5 hr at 37°C. Cells were washing in 50 μL of full media 5 min incubated at 37°C, and the wash was repeated three times. Images were taken at 580nm with Molecular Devices ImageXpress Microscope in live-cell compatible chamber supplied with 5% CO_2_.

### SNAP localization Assay using H1048 and BRM014

NCI-H1048 cells expressing SNAP-tagged proteins of interest were seeded in 12 well chamber slide (Ibidi #81201) at 15,000 cells per well in 200 μL of DMEM/F12 media (Corning) supplemented with 10% fetal bovine serum (Corning) and 1% penicillin/streptomycin (Gibco) 48 hours before the experiment. The cells were then treated with 3 μM of SNAP-cell TMR-STAR (NEB #S9105S) in 100 μL of full media and incubated for 0.5 hr at 37°C. Cells were washing in 100uL of full media 5 min incubated at 37°C, and the wash was repeated three times. Cells were fixed with 4% paraformaldehyde (Thermo #I28800) and permeabilized with 0.5% Triton-X in PBS. After removal of silicon chamber, the sample was mounted with Prolong Gold Antifade mounting solution with DAPI (Invitrogen #P36941). Images were taken at 580nm with Leica SP8 Laser Scanning Confocal Microscope.

### Cell viability and proliferation assay

Cell viability and proliferation rate were assessed through CellTiter-Glo (CTG) Luminescent cell viability assay (Promega). CTG working reagent was prepared according to manufacture protocol and was equilibrated to room temperature before experiment. In general, cells were seeded in white opaque 96-well plates (Corning) at 1,000 cells per well in 100 mL of media supplemented with 10 % fetal bovine serum (Corning) and 1% penicillin/streptomycin (Gibco) and allow to grow overnight before treatment. 100 uL of CTG working reagent was added to each well, and the plates were shaken at 450 rpm for 3 min for thorough mixing. The plates were incubated at room temperature for 10 min before luminesce reading was taken with Synergy H4 hybrid reader.

### SILAC cell-culture methods and proteomic sample preparation

SILAC labeling was performed by growing cells for at least five passages in lysine- and arginine-free SILAC medium (RPMI or DMEM, Invitrogen) supplemented with 10% dialyzed fetal calf serum (Gemini) and 1% Pen/Strep. “Light” and “heavy” media were supplemented with natural lysine and arginine (0.1 mg/mL) for “light”, and ^13^C-, ^15^N-labeled lysine and arginine (0.1 mg/mL) for “heavy”, respectively.

### Sample preparation and streptavidin enrichment

Quantitative proximity labeling studies with SILAC quantitative proteomics were performed with “heavy” and “light” labeled HEK293T lines. Quantitative proximity labeling studies with MDA-MB-231 were performed using LFQ. Cells, grown to 80-90% confluency in 10 cm cell-culture treated plates each, were incubated with DMSO alone (light cells) or PhotoPPI probe (15 μM, heavy cells) for 1 h in SILAC RPMI. After incubation, excess probe was removed by aspiration and supplanted with full media (10 mL x 10 min) for two cycles before replacement of media with PBS (5 mL). Cells were then irradiated using a Spectroline XL 1500 (UV, 365 nm) for 5 min, rinsed, scraped, and washed with cold PBS (1 ml x 4). The cells were pelleted and then lysed in RIPA lysis buffer (50 mM Tris, 150 mM NaCl, 1% Triton X-100, 0.5% deoxycholate, pH 7.4) supplemented with EDTA-free complete protease inhibitor (Roche) and 1 mM DTT, at 4 °C. After sonication, insoluble debris was cleared by centrifugation (17,000 *g*, 15 min). BCA assay was performed to normalize Probe and Vehicle protein concentrations to ∼1 mg/ml. Streptavidin agarose beads (50 μL slurry, Pierce) were washed twice with RIPA buffer, and each cell lysate was separately incubated with the beads with rotation overnight at 4 °C. The beads were subsequently washed five times with 0.5 mL of RIPA lysis buffer containing 1 mM DTT, combined together, then washed once with 1 mL of 1 M KCl, four times with 0.5 mL PBS, and two times with 2 M Urea in 25 mM ammonium bicarbonate. 500 μL of 6 M Urea in 50 mM ammonium bicarbonate was then added to the beads, and samples were reduced on resin by TCEP (10 mM final) with orbital shaking for 20 minutes at 65 °C. Samples were then alkylated by adding iodoacetamide (20 mM final), covered from the light and with orbital shaking, for 40 minutes at 37 °C. The streptavidin agarose beads were collected, washed once with 2 M Urea in 25 mM ammonium bicarbonate, and the buffer exchanged to 2 M Urea in 25 mM ammonium bicarbonate supplemented with 1 mM CaCl2. Enriched proteins were digested on-bead by the incubation of 2 μg sequencing grade trypsin overnight at 37 °C. Following trypsinization, supernatant was collected, acidified with HPLC grade formic acid (2% final, pH 2-3), and peptides were then desalted on ZipTip C18 tips (100 μL, Millipore), dried under vacuum, resuspended with LC-MS grade water (Sigma Aldrich), and then lyophilized. Lyophilized peptides were dissolved in LC-MS/MS Buffer (H2O with 0.1% formic acid, LC-MS grade, Sigma Aldrich) for proteomic analysis.

### LC-MS/MS Acquisition and Analysis

The proteomic methods reported are adopted from our previous reports^48^. LC-MS/MS analysis for proteomics samples was performed with an UltiMate 3000 RSLC nano System (Thermo Fisher Scientific) using an Acclaim PepMap RSLC C18 column (75 μm × 15 cm, 2 μm, 100 Å, Thermo Fisher Scientific) with an in-line Acclaim PepMap 100 C18 trap column (75 μm × 2 cm, 3 μm, 100 Å, Thermo Fisher Scientific) heated to 45 °C. The LC system was coupled to an Orbitrap Exploris 480 and Nanospray Flex Ion Source with stainless steel emitter tip (Thermo Fisher Scientific). Mobile phase A was composed of H_2_O supplemented with 0.1% formic acid, and mobile phase B was composed of CH_3_CN supplemented with 0.1% formic acid. The instrument was run at 0.3 μl min^−1^ with 2 h gradients. MS/MS spectra were collected for the entirety of the gradient using a data-dependent, 2-second cycle time setting with the following details: full MS scans were acquired at a resolution of 120,000, scan range of 380 *m*/*z* to 1,500 *m*/*z*, maximum IT of 25 ms, normalized AGC target of 300% and data collection in profile mode. MS2 scans were performed by high-energy collision dissociation (HCD) fragmentation with a resolution of 15,000, normalized AGC target of 50%, maximum IT of 50 ms, HCD collision energy of 30% and data collection in centroid mode. The isolation window for precursor ions was set to 1.6 *m*/*z*. Peptides with a charge state of 1, 7+ and unassigned were excluded, and dynamic exclusion was set to 40 seconds. The RF lens % was set to 40 with a spray voltage value of 2.0 kV and an ionization chamber temperature of 300 °C.

Data were processed using the SEQUEST HT search engine node within the Proteome Discoverer 3.0 software package. Data were searched using a concatenated target/decoy UniProt database of the human proteome with isoforms. Digest enzyme specificity was set to trypsin with up to two missed cleavages allowed, and peptide length was set to between 6 and 144 residues. Precursor mass range was set to 350–6500. Precursor mass tolerance was set to 10 ppm, and fragment mass tolerance was set to 0.02 Da. Up to 4 dynamic modifications were allowed per peptide, including heavy lysine (+8.0142), heavy arginine (+10.0083), oxidized methionine (+15.9949), N-terminal acetylation (+42.0106), N-terminal Met-loss (−131.0405) and N-terminal Met-loss + acetylation (−89.0299). Cysteine carboxyamidomethylation (+57.0215) was set as a static modification. A minimum of two peptides, with a minimum length of 6, was required for protein identification, and false discovery rate (FDR) was determined using Percolator with FDR rate set at 1%. Before quantification, chromatographic alignment was performed, with a maximum retention time difference of 10 min allowed, a mass tolerance of 10 ppm and a minimum signal/noise threshold of 5 required for feature mapping. SILAC ratios were determined using precursor-based quantification in a pairwise manner based on peak intensity without normalization or scaling using a maximum ratio of 20. For PhotoPPI studies using SILAC for quantitation, proteins considered as enriched interactors were detected and quantified in at least 2 biological replicates and exhibited a probe dependent median SILAC ratio greater than 4. MS files from LFQ samples were analyzed using the processing method parameters as indicated above with no dynamic modifications for heavy lysine or arginine. Ratios LFQ were determined using precursor-based quantification in a pairwise manner based on peak intensity without normalization or scaling.

STRING database version 12.0 was used to find known PPIs among the enriched proteins^49^. Physical networks from PPIs with experimental evidence from literature and databases (using interaction score of at least 0.150, text-mining removed) were selected in STRING database searches. Scatter plots showing the relationship between known PPIs and grouped ion intensity of enriched proteins were generated from the node degrees data from STRING analyses (Y-axis values) and the median grouped ion intensity (X-axis values) across biological replicates within the experiment at hand.

### Gene ontology and pathway analyses

Metascape analyses were performed on enriched proteins to identify significantly enriched gene sets and pathways. Gene sets from GO Molecular Functions, GO Biological Processes, Hallmark Gene Sets and KEGG Pathway were used for Metascape queries. Enrichment P-values from the Metascape analyses were used to generate plots to compare differences and similarities between sample types. In addition, some significantly enriched pathways were selected for detailed heatmaps showing abundance ratios of proteins belonging to the respective pathway. Heatmaps were generated using Morpheus software (Broad Institute) or Graph Pad Prism 8.

## Extended Data

**ED Figure 1:**
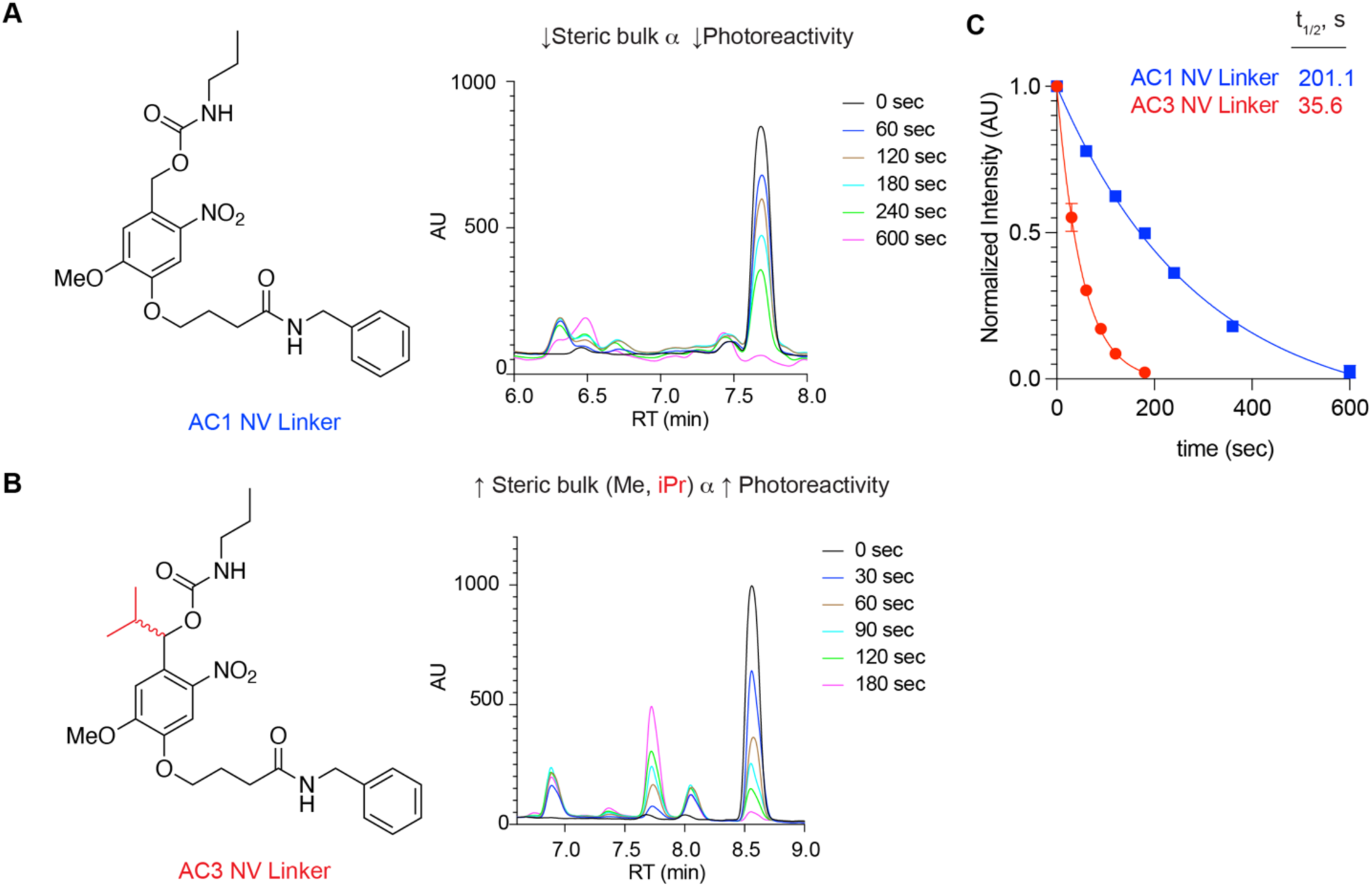
Photokinetic study using AC1 and AC3 nitroveratryl (NV) linkers. **A-B**) Chemical structure (left) and representative LC-MS traces (right) of AC1 (A) and AC3 (B) NV Linker after short pulses of 365 nm light. **C**) Kinetic plot of normalized intensity values for AC1 And AC3 NV model linker compounds in solution irradiated with pulses of UV light (365 nm) and monitored via LC-MS over indicated times. One-phase decay model was used to determine half-life for each compound using normalized absorbance unit (AU) intensity values at each time point; n = 3 biological replicates.

**ED Figure 2:**
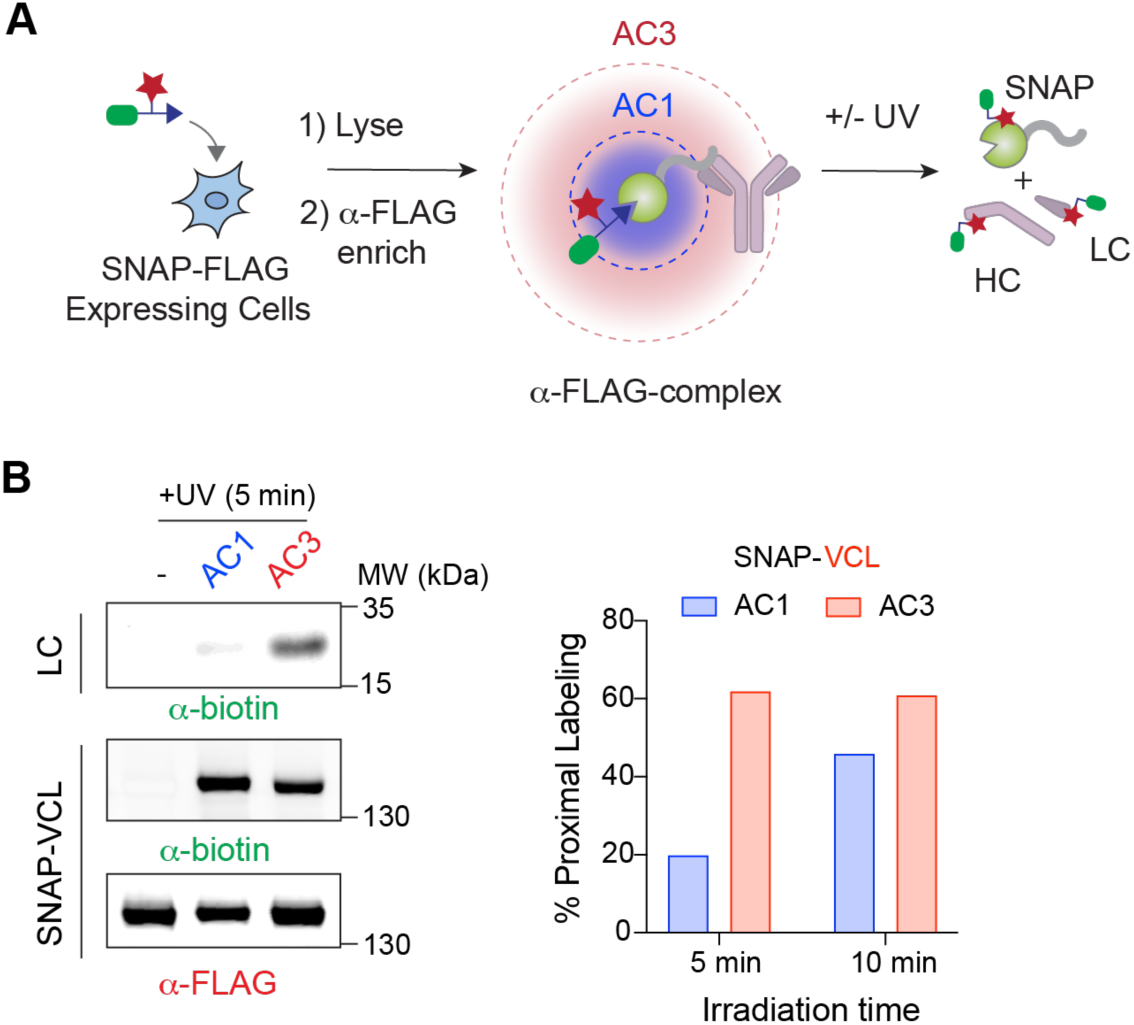
Comparative in-vitro SNAP complex photolabeling assay using siPROX. **A)** Schematic of *in vitro* FLAG-SNAP-Protein photolabeling assay. Cells or lysates are treated with photoprobe before enrichment of probe labeled construct and UV irradiation, resulting in labeling of the bait protein and α-FLAG antibody. Heavy Chain (HC) and Light Chain (LC) of antibody are shown. **B**) Left: Immunoblot for comparative in-vitro siPROX assay using SNAP-VCL. Biotin stained immunoblots are shown for the bait (SNAP-VCL) and PPI (Light chain, LC). Right: Bar plots showing quantification of in-vitro photolabeling data for SNAP-VCL; n = 2 biological replicates.

**ED Figure 3:**
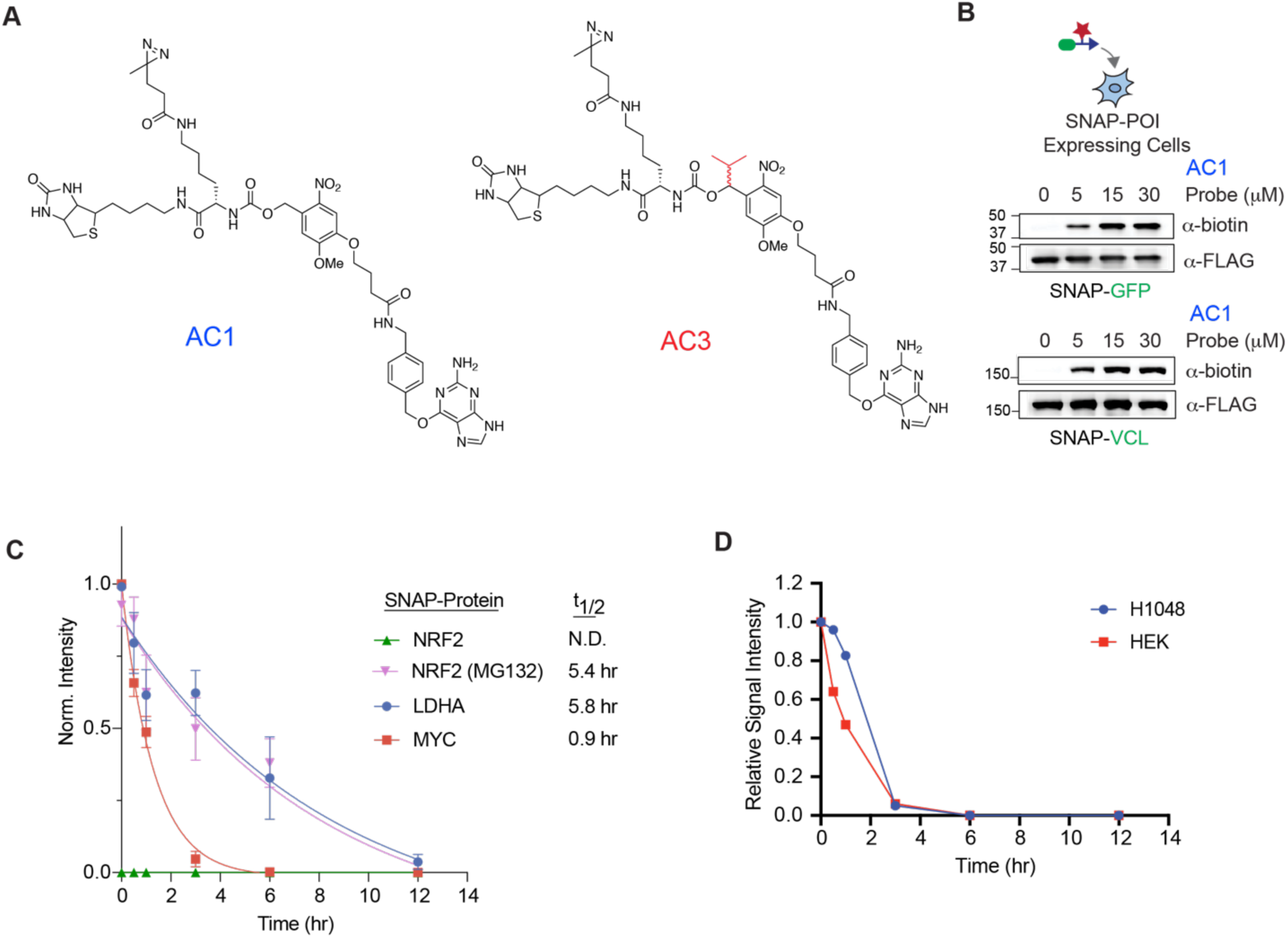
siPROX labeling in live cells. **A**) Chemical structures of siPROX photoproximity probes AC1 and AC3. **B**) Immunoblot (α-biotin, α-FLAG) from live cells treated with PhotoPPI probe AC1 for 1 h at the indicated doses (0-30 μM). Data shown is representative of at least n = 2 biological replicates. **C**) Quantification of probe labeled SNAP-Proteins over time in HEK293T cells stably expressing SNAP-NRF2, SNAP-LDHA or SNAP-MYC; n = 3. **D**) Quantification of probe labeled SNAP-MYC over time in HEK293T (red) and H1048 (blue) cells.

**ED Figure 4:**
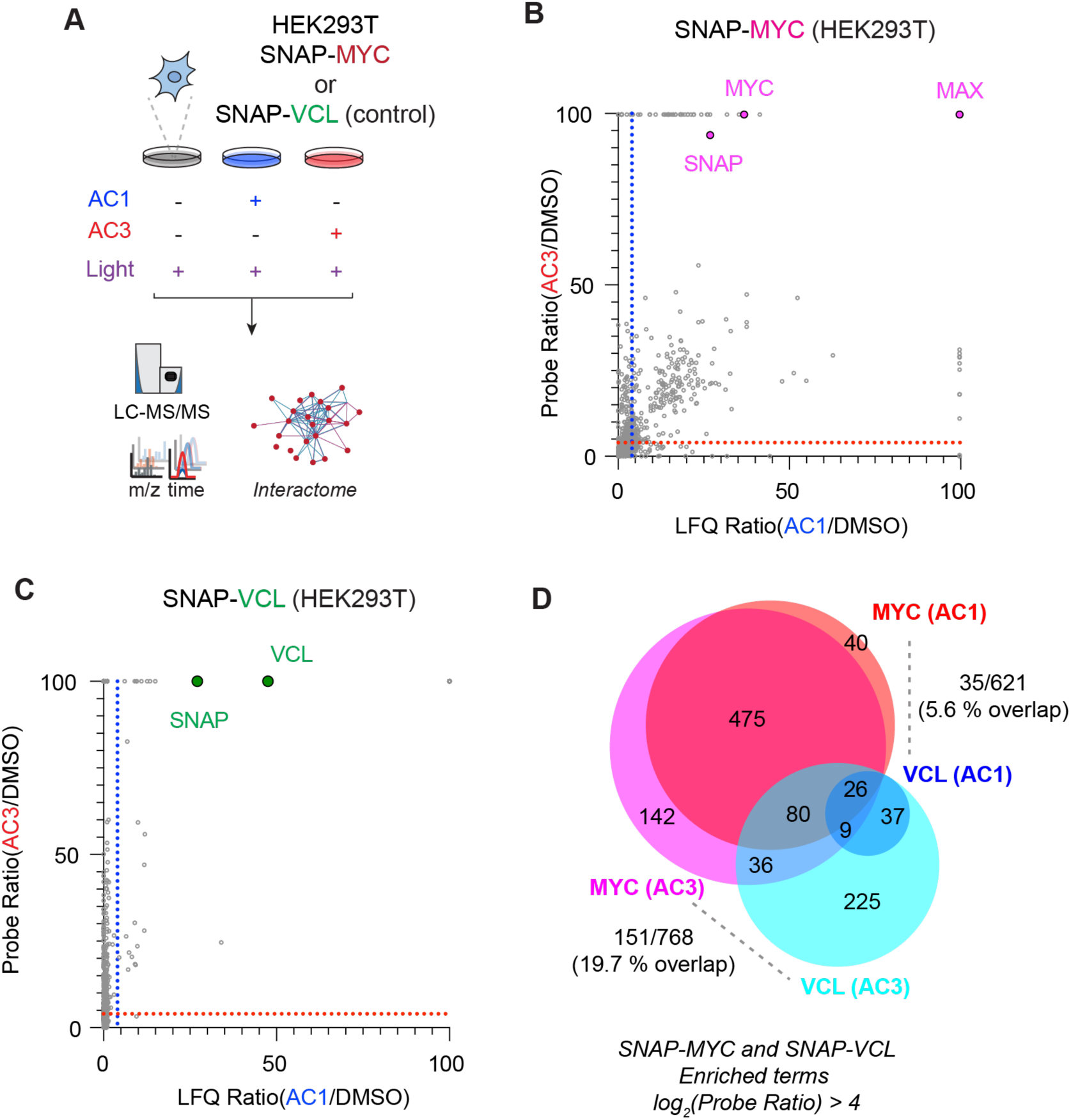
Drafting the MYC interactome using siPROX. **A**) Workflow for SNAP-MYC interactome profiling using relative label-free quantification (LFQ) measurements of probe/no-probe. SNAP-VCL expressing HEK293T cells were used as control **B-C**) Correlation plots showing AC1 and AC3 probe LFQ ratios (Probe/no-probe) for SNAP-MYC (B) and SNAP-VCL (C). **D**) Venn diagrams showing shared and unique enriched AC1 and AC3 targets from the profiles shown in B-C. Data is representative of n = 4 biological replicates.

**ED Figure 5:**
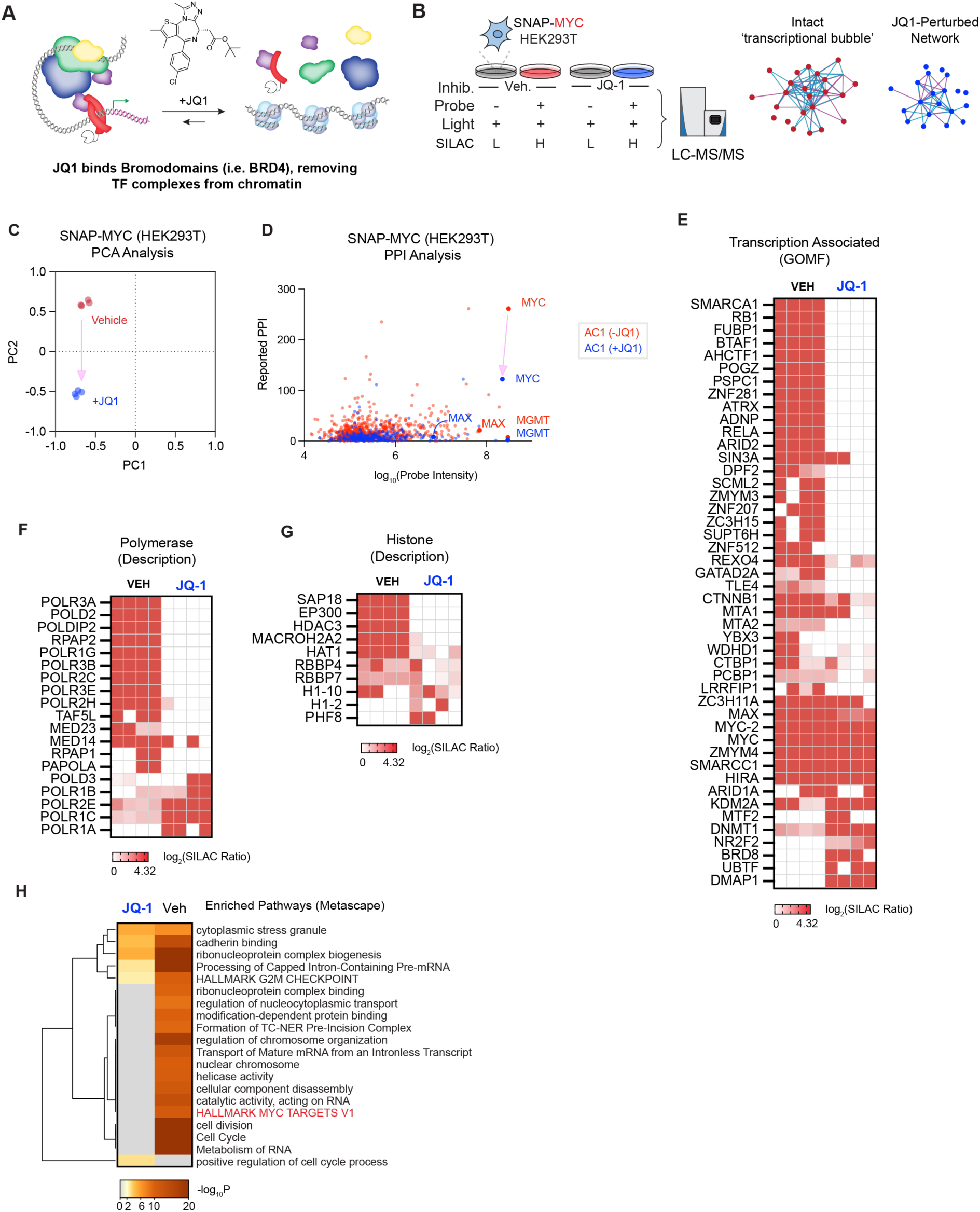
Global MYC interactome analyses following chronic bromodomain inhibition. **A)** Graphical depiction of JQ-1-mediated disruption of bromodomains causing removal of transcription initiating complexes from chromatin. **B**) Experimental outline for profiling the MYC interactome in the presence of vehicle or JQ-1 to identify novel interactome rewiring events. SNAP-MYC cells are pretreated with JQ-1 for 24 h prior to siPROX profiling under standard conditions. **C)** Principal component analysis (PCA) of basal and JQ-1 perturbed interactomes. **D**) Scatter plot depicting the relationship between the grouped ion-intensity and the number of STRING database-annotated protein-protein interactions (PPIs) for each enriched protein (ratio > 4) in Vehicle (Red) and JQ-1 (Blue) treated cells using AC1 probe; n = 4 biological replicates. **E-G**) Heatmaps of enriched proteins from Vehicle or JQ-1 treated cells; keywords used: ‘transcription’ in GOMF (E), ‘polymerase’ (F), or ‘histone’ (G). **H**) Metascape analysis of AC1 derived SNAP-MYC interactomes under vehicle or drug-perturbed conditions.

**ED Figure 6:**
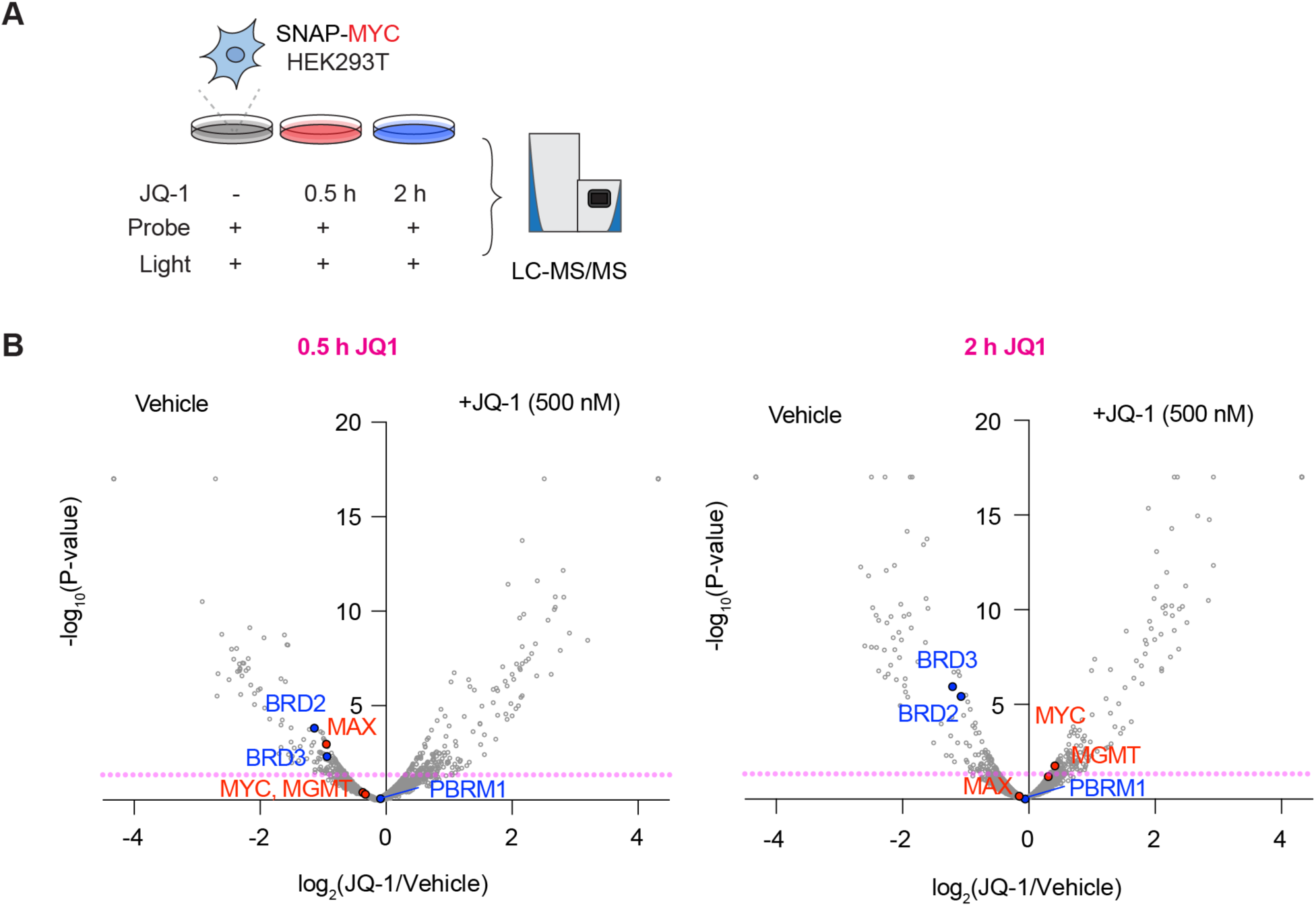
Rapid complex dynamics measured after short treatments of JQ-1. **A**) Experimental model for rapid dynamic profiling of JQ-1 interactome effects. SNAP-MYC cells are pretreated with JQ-1 for 30 min and 2 h prior to siPROX profiling under standard conditions. B) Volcano plots of log_2_(JQ1/Vehicle) plotted against -log_10_(p-value) at 30 min (left) and 2 h (right) of JQ-1. Proteins that are significantly perturbed by JQ-1 contained a *p*-value less than 0.05 at each time point; data is representative of n = 2 biological replicates at each time point.

**ED Figure 7:**
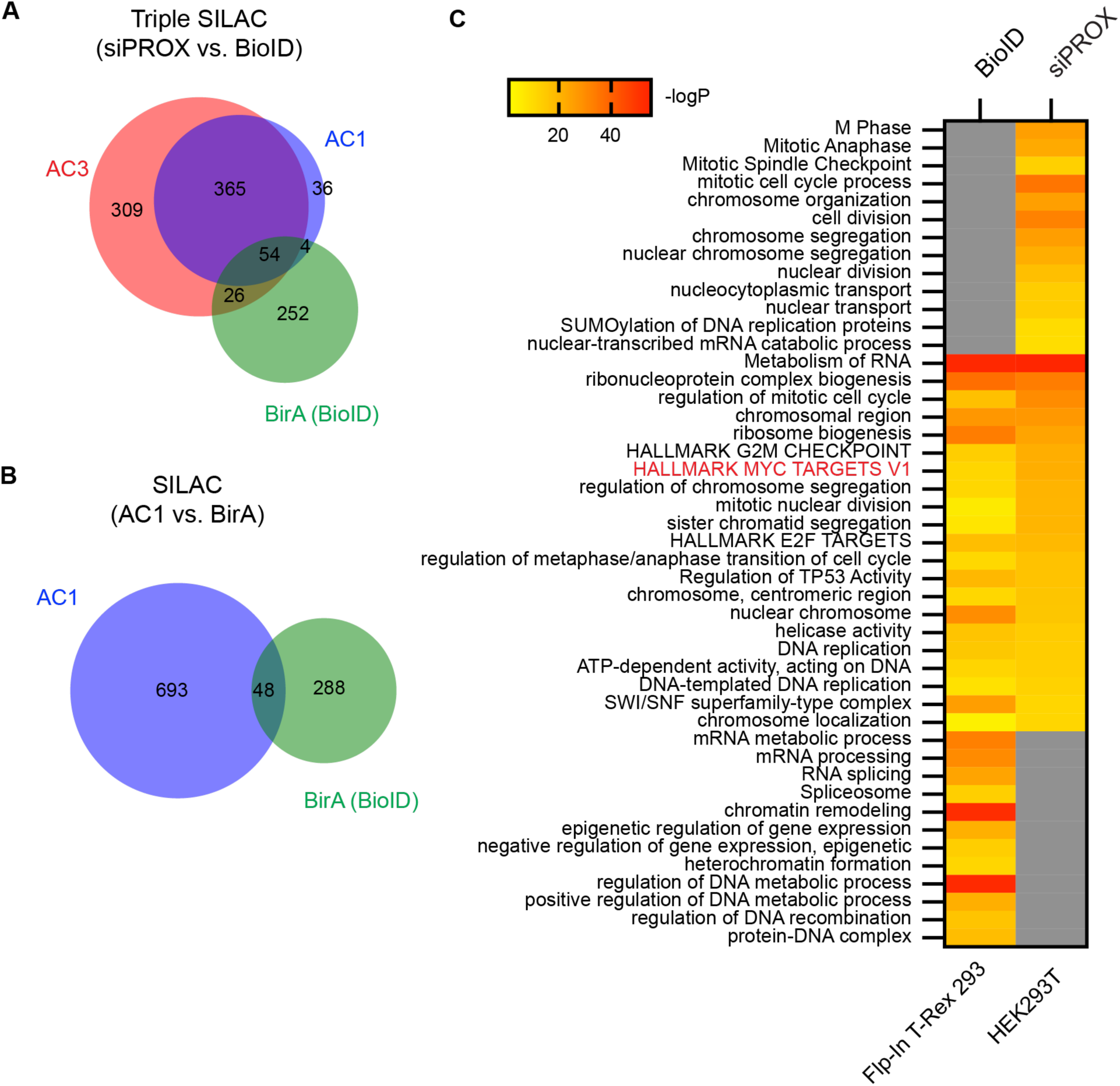
Comparison of MYC interactomes derived from siPROX and BioID. **A**) Venn diagram showing shared and unique MYC PPIs as identified by siPROX or BioID. Comparative analysis was conducted using gene symbols. **B**) Venn diagram showing shared and unique MYC PPIs as identified by siPROX (AC1, Main Figure 1) or BioID (BirA). **C**) Heatmap showing results from Metascape analysis of enriched terms from BioID or siPROX profiling using input terms from the analysis in B.

**ED Figure 8:**
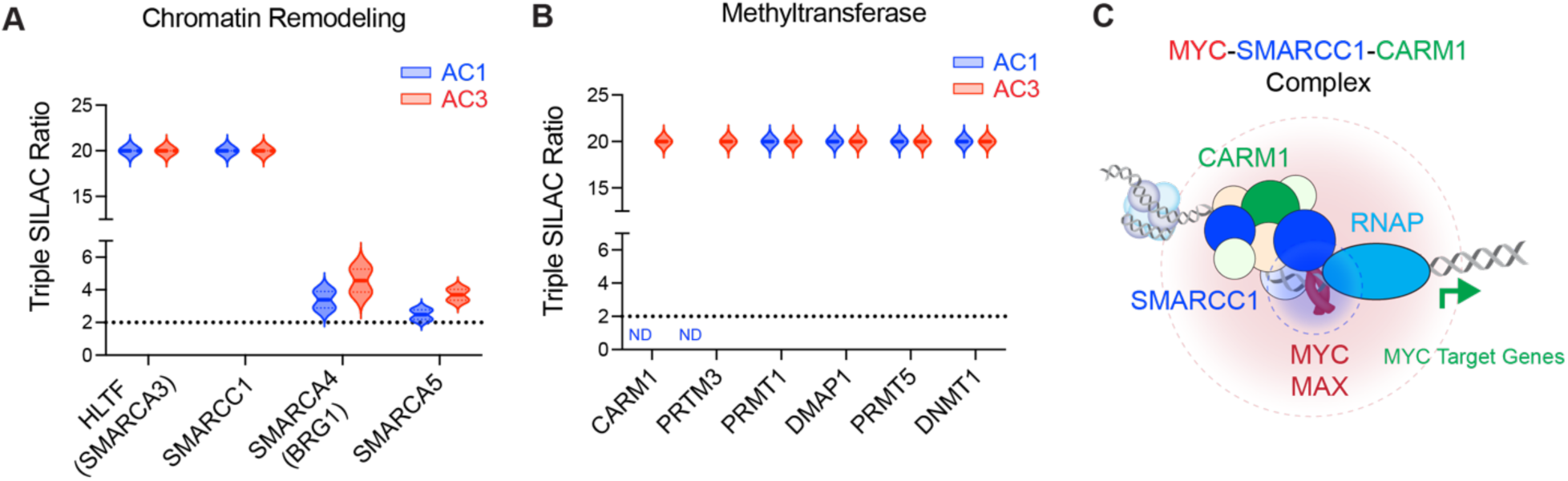
siPROX mapping of the MYC interactome in HEK293T cells reveals a spatially resolved methyltransferase-BAF-MYC transcriptional complex. **A-B**) Box and violin plots showing the Triple SILAC derived ratios for SMARC chromatin remodelers (A) and methyltransferases (B) enriched by siPROX; n = 2 biological replicates for each probe. **C**) Graphical depiction of the MYC-SMARCC1-CARM1 complex showing an indirect relationship between the MYC-MAX complex and CARM1 as determined from siPROX profiling.

**ED Figure 9:**
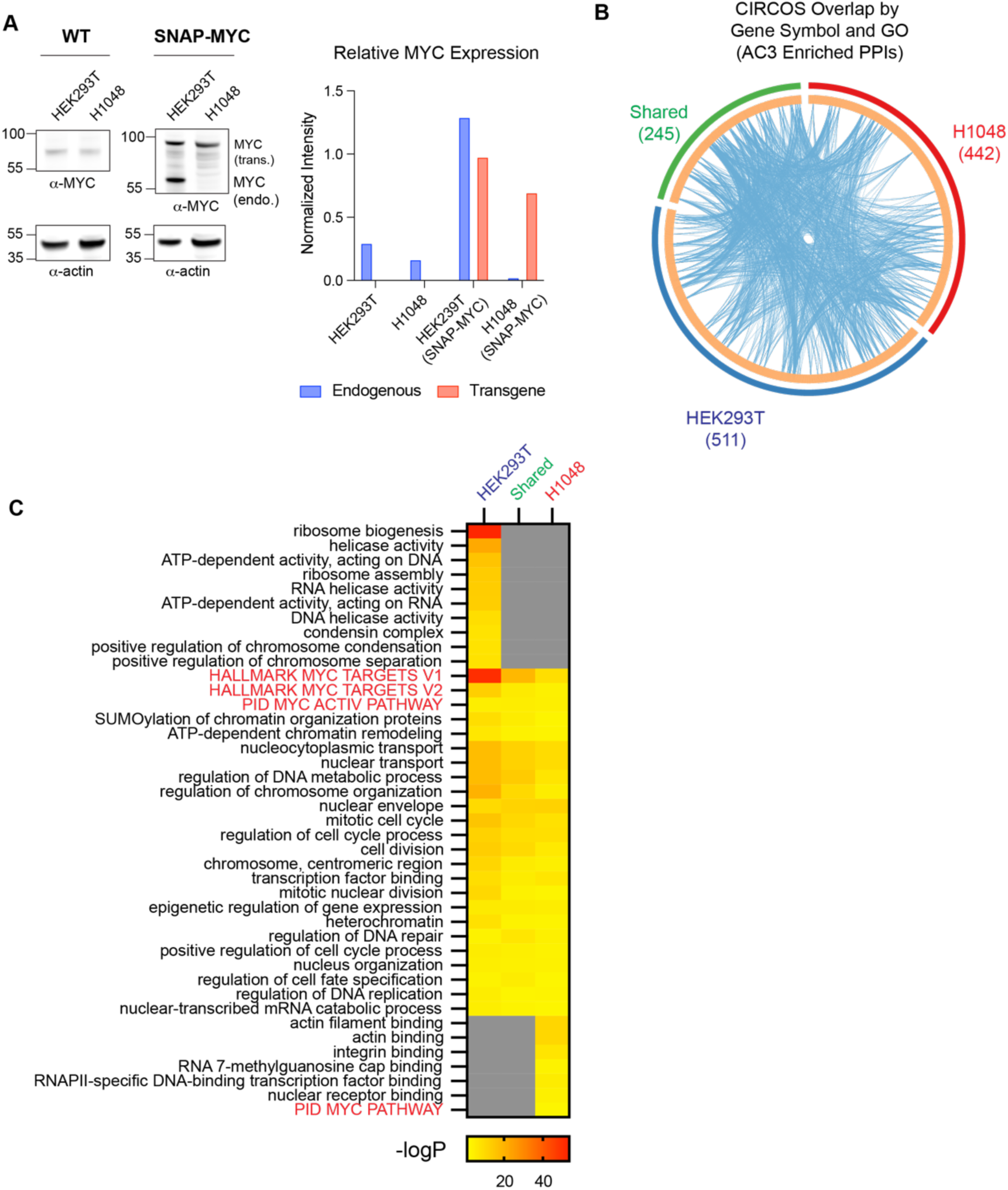
Overexpression of MYC in disparate cell lines alters MYC interactomes revealing common and cell-specific pathways. **A**) Relative MYC levels in HEK293T and H1048 cells overexpressing SNAP-MYC. Left: α-MYC and α-Actin immunoblots for WT and SNAP-MYC expressing cells. Right: Bar plot quantification of MYC expression levels normalized to actin; data is representative of n = 2 biological replicates. **B**) Circos plot constructed with gene symbols and GO annotations for AC3-derived MYC interactomes in HEK2923T and H1048 cell overexpressing SNAP-MYC. **D**) Metascape analysis of enriched pathways derived from siPROX profiling of SNAP-MYC expressing HEK293T and H1048 cells.

**ED Figure 10:**
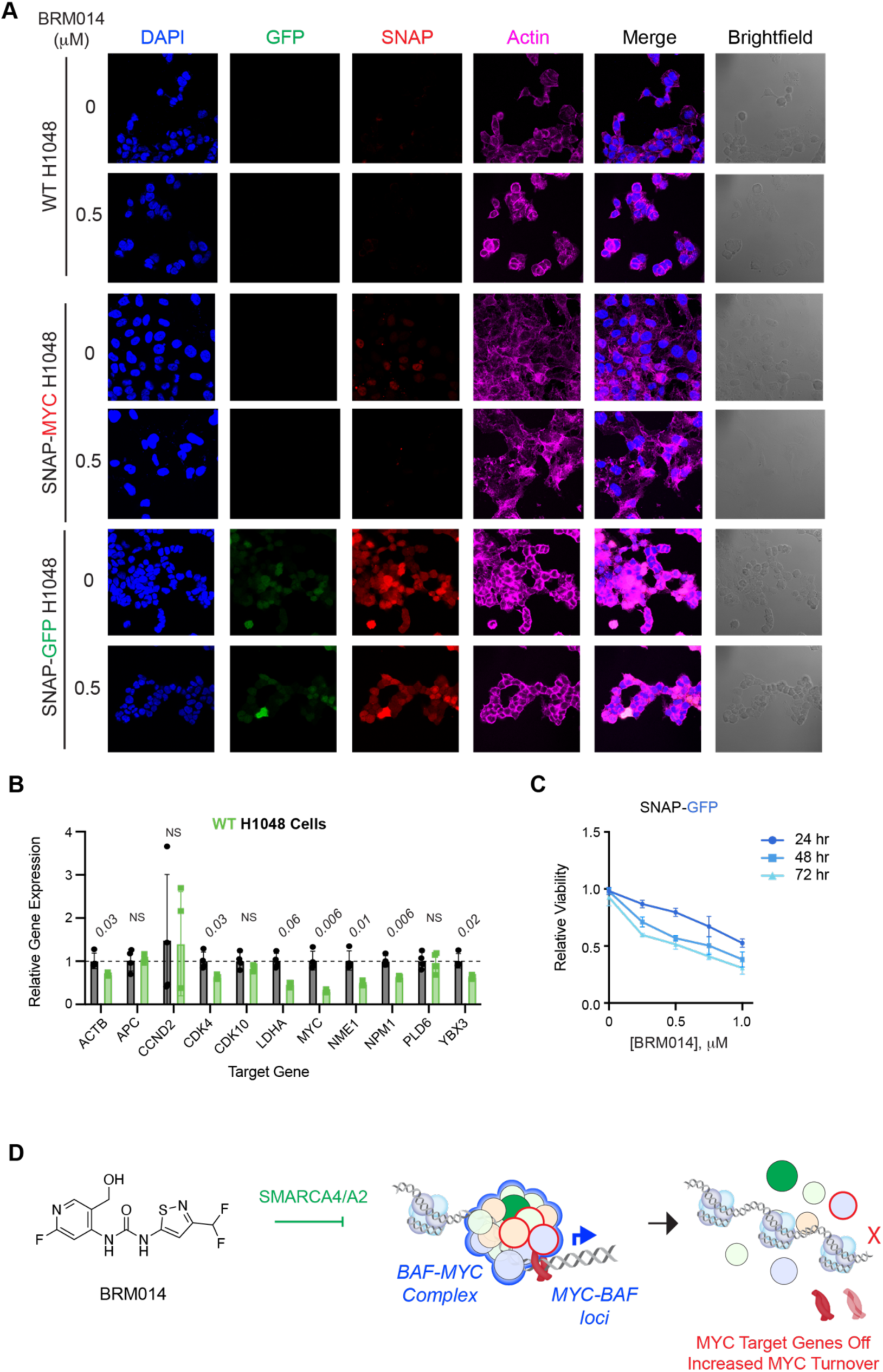
BAF complex inhibition reduces MYC-dependent signaling and reduces proliferation in models of MYC-driven cancer. **A)** Confocal microscopy of SNAP-MYC, SNAP-GFP and WT H1048 cells treated with BRM014 inhibitor (500 nM, 18 h) pretreatment. **B)** Relative qPCR analysis on MYC target genes in response to BRM014 treatment in WT H1048 cells. **C**) Representative confocal images of WT, SNAP-MYC and -GFP H1048 cells treated with BRM014 inhibitor (500 nM, 18 h). **D**) Graphical depiction of MYC-BAF complex inhibition by BRM014 targeting of SMARCA4/A2, which leads to increased MYC destabilization and turnover.

## Acknowledgements

We thank S. Ahmadiantehrani for her valuable suggestions during the writing process, figure drafting and proofreading assistance. This work was supported by NIH Multidisciplinary Training Grant in Cancer Research (MTCR) T32-CA09594 (to A.C.), and R01-GM145852 (to R.E.M), the Komen Career Catalyst Research CCR 21663985 and Alfred P. Sloan FG–2020–12839 (to R.E.M.).

